# Artificial Cells Evidence Apical Compressive Forces Building up During Neuroepithelial Organoid Early Development

**DOI:** 10.1101/2025.06.13.659497

**Authors:** Hassan Kanso, Stefania Di Cio, Ruth Rose, Isabel Palacios, Julien E. Gautrot

## Abstract

During early stages of development of cerebral organoids, budding neuroepithelia display striking changes in size and morphology, occurring very rapidly. Whilst mechanical forces mediated by cadherin-cadherin junctions are known to control the assembly, maturation and stability of epithelia, little is known of the mechanical context associated with neuroepithelial organoid development. In this report, we demonstrate a rapid translocation of YAP to budding neuroepithelial apical junctions, suggesting the build-up of strong compressive forces early on in their development. To study the mechanics of budding rosettes, we designed oil microdroplets stabilised by protein nanosheets displaying cadherin receptors, able to engage with receptors presented by neighbouring neuroepithelial cells, to integrate into embryoid bodies and developing organoids. The resulting artificial cells are able to sustain the formation of mature junctions with neighbouring cells and lead to the recruitment of tight junction maturation proteins such as ZO1. During early budding of neuroepithelial rosettes, artificial cells are found to be rapidly expelled from the developing organoids, further evidencing apical compressive forces. These forces are not opposed by sufficiently strong shear forces from neighbouring cells, or adhesive forces maintaining anchorage to the apical junction, to induce deformation of artificial cells.

## 1. Introduction

Mechanical forces continuously shape tissue development and the correct regulation of tissue architecture and function [1–3]. During gastrulation, apical contractility of the actomyosin cytoskeleton guide invagination of the blastula to form the three germ layers [4]. During neurulation, cell-generated mechanical forces within the flat epithelium of the neural plate reshape and transform this tissue into the neural tube [5]. Throughout neuroepithelial development and neuro-morphogenesis, the mechanical forces generated during neural tube formation [5–7], the role of mechanical forces at cell-cell junctions in the shaping of epithelia [1,8], and the rapid morphological changes and folding occurring [9–11] underpin a critical role for complex biomechanical cues.

Organoids offer unique opportunities to investigate some of the processes controlling tissue development in a human context. Since the first report of intestinal organoids [12], a broad range of systems have been proposed, including kidney [13], lung [14], skin [15], bone marrow [16], eye [17] and cerebral organoids [18,19], amongst many others. Some of these models can be readily embedded in engineered matrices other than Matrigel, which allowed to demonstrate the importance of the mechanics of the organoid microenvironment in the regulation of cell differentiation and patterning of the developing tissue. For example, hydrogels with controlled degradation enable the occurrence and positioning of intestinal crypt development [20,21], whilst lung organoid maturation is regulated by hydrogel mechanics [22].

Probing mechanical forces in organoids is particularly challenging as it is difficult to access contractile or compressive events taking place deep in these cellular assemblies. Beyond direct contact probing techniques, such as force probe microscopy, which are restricted to probing surface mechanics, with limited extrapolation to core mechanics [23], optical and magnetic tweezer-based methods allow the probing of deeper mechanics in cellular assemblies [24]. More recently, microdroplet force sensors have attracted attention as they enable reaching the deeper mechanics of cell assemblies and even embryos [25,26]. For example, this enabled the identification of fluid flow and jamming during tissue morphogenesis and embryogenesis [25] and provided evidence for anisotropic compressive forces shaping the development of the murine incisor [27].

YAP and TAZ are transcriptional co-activators playing an important role in the regulation of cell proliferation and differentiation [28,29], the correct orchestration of organoid development [20,22] and have a significant impact on brain development at multiple stages, from neural tube formation to maturation of the cerebral cortex and cerebellum [30]. Deletion of YAP leads to abnormal neural tube closure [31], its silencing reduces neural crest cell migration [32] and the balance of YAP activation regulates progenitor proliferation in the developing neural tube and ultimate neuron density [33]. In turn, during later cortical and cerebellar development, YAP and TAZ were shown to regulate the shape, architecture and neuronal density [34–36]. Similarly, conditional knock out of YAP led to impaired retinal progenitor cell proliferation and cell cycle progression, leading to retina developmental defects [37] and, in optic neuroepithelial development, the drosophila YAP homologue Yki controls the cell cycle arrest preceding conversion to neuroblasts [38].

The translocation of YAP from the cytosol to the nucleus has been shown to constitute an important process mediating mechanosensing and the response of cells to substrate mechanics, cyclic strain and extra-cellular matrix geometry [28,29], mediated by the Ras family GTPase RAP2 [39]. YAP translocation and associated mechanosensing itself is regulated by and regulates the formation of focal adhesion and the assembly and contractility of the actin cytoskeleton [40,41], including in 3D microenvironments [42]. In turn, these processes were found to impact a broad range of cell phenotypes, including the differentiation of mesenchymal stem cells into multiple lineages [28,43], their remodelling of the pericellular matrix [42], myogenesis [44] and cardiomyocyte proliferation [45], hepatocyte function [46] and the balance of expansion and differentiation in the interfollicular epidermis [47,48]. In addition, YAP-regulated mechanosensing is not restricted to matrix-mediated processes and equally plays a critical role in the regulation of force sensing by cadherins at adherens junctions [49,50].

Although the importance of mechanosensing and YAP-mediated signalling and the Hippo pathway are well established in the development of organoids, including intestinal [51,52], liver [53,54] and lung organoids [55], the temporal regulation of YAP translocation during neuroepithelial organoid development has not been studied. However, mechanical stimulation of neuroepithelial organoids using low intensity ultrasounds led to enhanced growth in a YAP-dependent manner (although investigated via direct knock down of YAP) and enhanced the maturation of neural lineages in vitro and upon implantation in mice hosts [56]. Stimulated microgravity caused impaired development (reduced proliferation and abnormal architecture) of neuroepithelial organoids, associated with reduced Hippo pathway activation, and rescued by YAP activation via MST1/2 inhibition [57]. In contrast, confinement in microgrooves led to inhibition of YAP and reduction in lumen formation and FOX2A patterning [58]. Similarly, bioreactor culture and differentiation of hPSC-derived primitive neural stem cells led to enhanced neuronal lineage maturation compared to 2D cultures, with associated YAP1 cytoplasmic translocation [59].

Hence, we proposed that mapping the evolution of YAP localisation during early stages of the formation of neuroepithelial organoids could shed light on the evolution of their mechanical context. To probe associated mechanical forces developing during neuroepithelial maturation, we then applied microdroplet force sensors. However, to avoid having to inject microdroplets into organoids, resulting in their dissociation and damage, and to enable direct engagement of cell-cell receptors, such as cadherins, we proposed to develop microdroplet-based artificial cells presenting defined cadherin receptors and enabling embedding into embryoid bodies prior to neuroectoderm induction and differentiation. In this report, we demonstrate the assembly of such artificial cells, their embedding in embryoid bodies and their rapid expulsion from budding neuroepithelia, evidencing the build-up of compressive forces occurring during early rosette formation, associated with YAP translocation.

## 2. Results and discussion

### 2.1. The mechanotransducer YAP translocates upon initiation of neuroepithelial development

Neuroepithelial organoids were generated via neuroectoderm induction of embryoid bodies, followed by neuroepithelial differentiation and maturation (Figure 1A) [18]. During this process, the initially smooth spherical embryoid bodies displayed a gradual roughened surface associated with symmetry breaking occurring during the onset of multiple neuroepithelia forming in each organoid, followed by expansion of their size and overall volume of the organoid (Figure 1A and Supplementary Videos S1-2) [19]. Immunostaining revealed that neuroepithelial rosette budding occurred from day 1 of differentiation, followed by lumen formation, enlargement and maturation (Figure 1B), in agreement with literature observations [19]. During this time frame, YAP was found to be initially nuclear and gradually translocated to the cytosol, with relatively little nuclear fraction remaining by day 9 (Figure 1C-D). In addition, we found that YAP localisation at this end time point was primarily at the apical junction of developing neuroepithelia.

**Figure 1.**
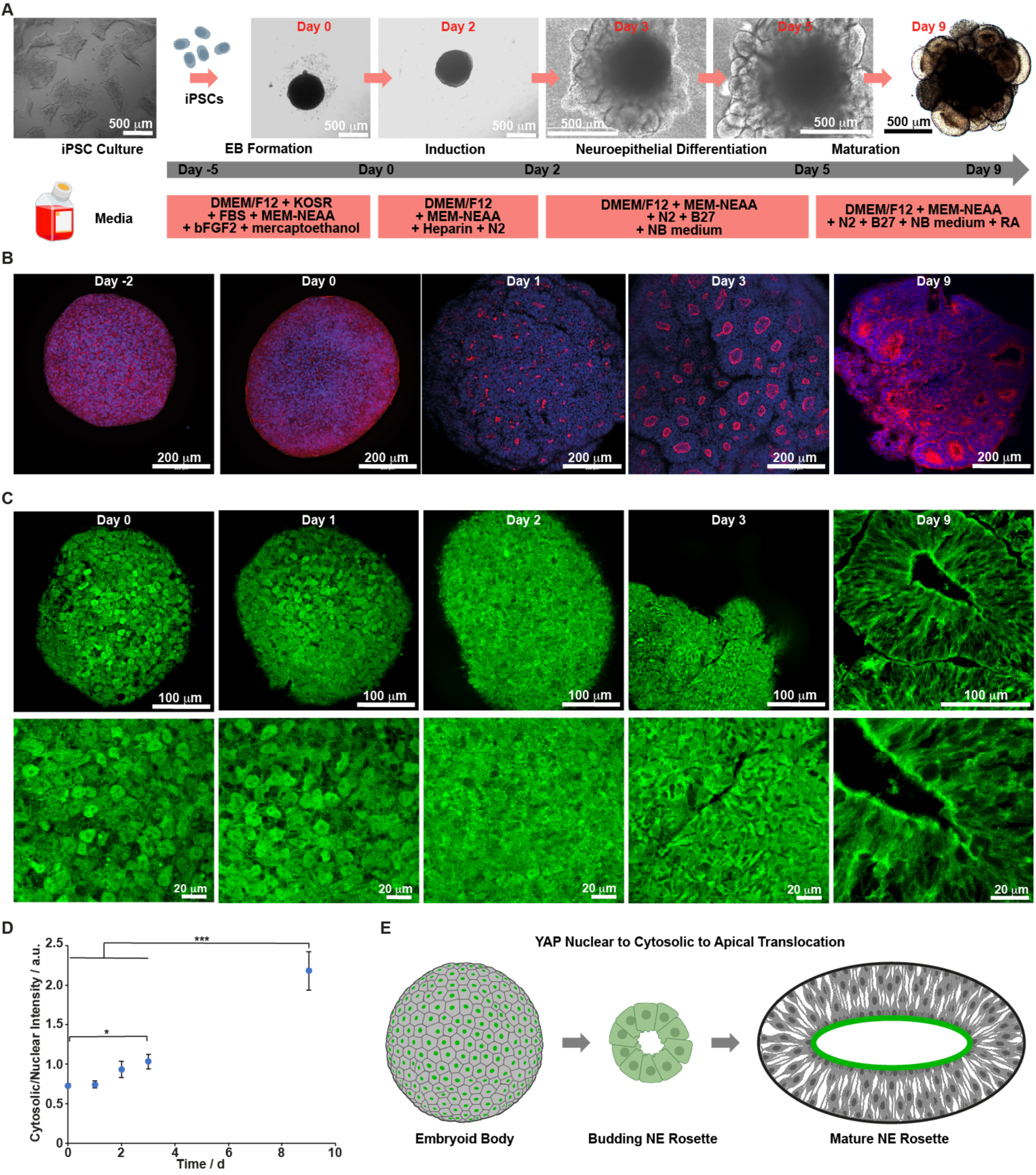
YAP Cytosolic translocation is initiated early during neuroepithelial rosette development and then accumulates at apical junctions. A. Schematic representation of the differentiation protocol applied to generate neuroepithelial organoids and representative bright field microscopy images of developing organoids at different stages. B. Fluorescence microscopy images (max projection of confocal stacks) of developing organoids. Blue, DAPI; Red, ZO1; scale bar: 200 μm. C. Fluorescence microscopy imaging of YAP localisation (single optical sections taken from confocal stacks) in developing organoids at indicated time points. Day 9 are images of histological sections, and all other images are taken from confocal stacks obtained for full embryoid bodies/organoids. Green, YAP; scale bar: 100 μm. D. Quantification of cytosolic/nuclear fluorescence intensity measured in corresponding images. N = 3 ± SEM; *, p < 0.05; ***, p < 0.001. E. Schematic representation of the proposed translocation of YAP (indicated in green) occurring during neuroepithelial rosette formation and maturation.

Although apical localisation of YAP, at tight junctions formed by the developing neuroepithelial organoids was not previously reported [57–59], it is in good agreement with its localisation at apical junctions formed by radial glial cells in the developing ventricular zone [34]. This is also in good agreement with its reported localisation at the apico-basal junctions of retinal section at E15.5 [37]. YAP apical junction distribution was also reported in colonic crypt cells, where its accumulation at adherens and tight junctions was regulated by RAB11A [60]. Therefore, our observations indicate that YAP remains mainly nuclear following neuroectodermal induction of embryoid bodies, and then gradually translocates to the cytosol during early onset of neuroepithelial rosette budding, following by its accumulation at apical junctions during neuroepithelial rosette maturation (Figure 1E).

Considering the importance of YAP cytoplasmic translocation typically observed during cell spreading, response to substrate mechanics [28,46], including in 3D matrices [42], and as a result of mechanical strain and tugging forces generated at cell-cell junctions [49,50], our observations suggest that mechanical forces build up in neuroepithelial rosettes, even at early stages of development. Indeed, mechanical regulation of YAP translocation was observed in other contexts. In drosophila, during oogenesis and development of the egg chamber, Yki accumulated at apical junctions of columnar cells and in the nuclei of stretch cells [61]. Similarly, in MDCK cells and keratinocytes, high cell densities resulted in YAP translocation to the cytosol, in a process regulated by Merlin and the contractile circumferential apical actin belt [62,63]. Therefore, we proposed that YAP translocation in early stages of neuroepithelial development underpinned the build-up of mechanical forces.

### 2.2. Protein engineering to design artificial cells

Microdroplets stabilised by protein nanosheets have been demonstrated to enable cell adhesion, spreading and the maintenance of cell phenotype [64,65]. In these systems, the control of interfacial shear mechanics, viscoelasticity and toughness, in addition to the presentation of suitable adhesive ligands, enabled corresponding interfaces to resist cell mediated contractile forces, allowing cell spreading and the downstream integrin-mediated regulation of cell proliferation and stemness retention [66,67]. In turn, this enabled the long-term maintenance of mesenchymal stem cells [68], the control of their differentiation [69,70], the proliferation of induced pluripotent stem cells and direct induction of differentiation to cardiomyocytes, without further processing [71], and the in vitro expansion of haematopoietic stem cells [72]. Here, we proposed the design of microdroplet-based artificial cells, stabilised by protein nanosheets, controlling interfacial mechanics (mimicking cortical mechanics), and presenting cadherins to mimic cell-cell adhesive cues and allow the direct capture and integration of microdroplets by cellular assemblies (Figure 2A). To build artificial cells with controlled architecture and mechanics, we selected albumin as a scaffolding amphiphilic protein for the stabilisation of microdroplets, allowing a broad range of interfacial mechanics to be achieved [73], and introduced single biotin residues recombinantly via an Avi tag (Supplementary Figures S1-2). Subsequent binding of streptavidin and biotinylated Protein G was proposed to enable the capture of Fc fusion cadherins, such as E- and N-cadherins. Defined biotinylation was selected to avoid crosslinking within the protein layer. To ensure that the tag selected was accessible for subsequent binding, without compromising the self-assembly of albumin to the surface of droplets, we selected the N-terminus of bovine serum albumin (BSA), which was shown by molecular dynamics simulations to remain dangling in the bulk aqueous phase following adsorption of this protein to hydrophobic surfaces such as graphite [74]. This is in agreement with the position of the Sudlow site II of BSA, a hydrophobic binding pocket present at the C-terminus, proposed to unfold upon adsorption at hydrophobic interfaces [74,75]. To validate that the small modification made to BSA (Avi tag with glycine bridge) would not impact on the globular assembly of BSA, we carried out structure prediction using AlphaFold [76], confirming an excellent retention of the bundled α-helices of native BSA and the accessibility of the biotin residue (Supplementary Figure S1).

**Figure 2.**
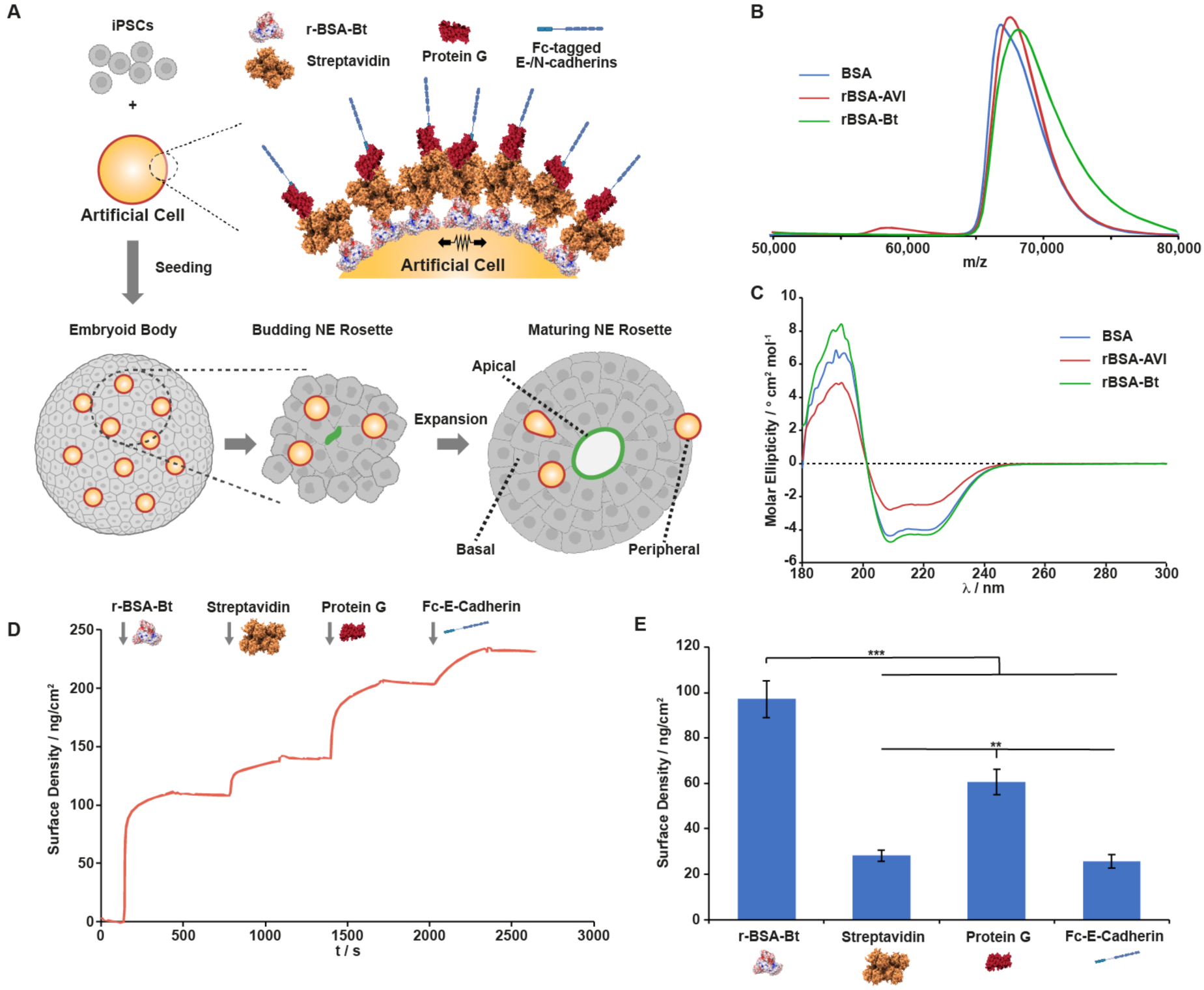
Engineering of recombinant protein nanosheet-stabilised artificial cells. A. Schematic representation of artificial cells generated through the self-assembly of protein nanosheets at the surface of oil microdroplets, and their incorporation into embryoid bodies, to probe the internal forces generated during the formation of neuroepithelial rosettes. B. Mass spectrometry traces of native BSA, rBSA-Avi and rBSA-Bt. C. Circular dichroism traces recorded in 5 mM phosphate buffer at 1 mg/mL for native BSA, rBSA-Avi and rBSA-Bt. D. Sequential adsorption of rBSA-Bt (1 mg/mL) at a fluorophilic monolayer, followed by binding of streptavidin (50 μg/mL), biotinylated protein G (100 μg/mL) and Fc-E-cadherin (25 μg/mL), in PBS, monitored by SPR at a flow rate of 10 μl/min. E. Corresponding quantification of protein densities measured by SPR. N = 3 ± SEM; **, p < 0.01; ***, p < 0.001.

To produce relatively high concentrations of the selected recombinant BSA (rBSA), we opted for yeast expression in *P. pastoris*, as this host has been demonstrated to support the expression of albumins in high yields and scales, with excellent folding and structure retention [77]. We selected the PIC9K vector for expression as this features an AOX1 promoter and α-factor secretion signal, allowing the release of the produced protein in the culture medium [78]. Successful transformants were identified at high concentrations of G418 to ensure sufficient copies of the rBSA plasmid, normally resulting in higher protein yields. Successful transformation and supernatant expression/secretion was achieved in good yields, in agreement with the literature for albumins [77], followed by purification of the resulting rBSA-Avi via Ni-NTA column (Supplementary Figures S3-5). The resulting rBSA-Avi was then converted into rBSA-Bt using recombinant BirA, with biotin conversion levels measured > 90%.

The resulting proteins were characterised by mass spectrometry, confirming the increase in molecular weight of rBSA-Avi and rBSA-Bt, compared to native BSA, and implying comparable glycosylation levels compared to this protein (Figure 2B). In addition, circular dichroism confirmed the high α-helix content typical of BSA, which adopts a highly globular conformation with tightly bundled helices (Figure 2C) [79]. Finally, the dual activity of the rBSA-Bt generated was confirmed by surface plasmon resonance (SPR). To model the fluorinated oil surface targeted to form microdroplets, fluorinated thiol monolayers were generated at the surface of SPR chips and rBSA-Bt was directly adsorbed to these interfaces, in PBS (Figure 2D). In agreement with the amphiphilic properties of globular albumins, rBSA-Bt rapidly adsorbed to fluorinated monolayers, to surface densities of 97 ± 8 ng/cm^2^ (Figure 2E). This is in good agreement with BSA adsorption levels reported for fluorinated monolayers, near 120 ng/cm^2^ [65], indicating the formation of monolayers with moderate packing. Upon subsequent adsorption of streptavidin, biotinylated Protein G and Fc-tagged E-Cadherin, successive rapid adsorption profiles were observed, with surface densities of 28 ± 2, 61 ± 6 and 26 ± 3 ng/cm^2^, respectively. Hence, our data confirms the formation of monolayers of cadherins captured at the surface of protein nanosheets, with moderately dense packing.

### 2.3. Protein nanosheet assembly to control artificial cell formation

The adsorption of protein nanosheets at fluorinated oil interfaces was investigated next. To quantify protein adsorption in situ and the mechanical properties of the resulting assemblies, interfacial shear rheology was carried out (Figure 3A). Upon injection of BSA or rBSA-Bt in the aqueous phase, the interfacial shear storage and loss moduli were found to increase from 0.1-1 mN/m to 4-5 mN/m and 1-2 mN/m, respectively, with the increasing gap between these two moduli and associated reduction in interfacial tan δ, confirming the formation of viscoelastic assemblies (Supplementary Figure S6). This was further confirmed by analysis of the frequency sweeps obtained after assemblies of albumin nanosheets, displaying modest changes in interfacial shear mechanics, for both proteins, over a broad frequency range (Figure 3B), in agreement with interfacial shear rheology previously reported for globular albumins [80,81]. Interestingly, upon introduction of streptavidin (after washing of excess rBSA-Bt), no further increase in modulus was observed (Figure 3C and Supplementary Figure S6), implying that the network mechanics is primarily dominated by the underlying recombinant albumin nanosheet formed, and that streptavidin binding did not induce further crosslinking in this assembly. Although this is somewhat surprising as streptavidin tetramers can bind 4 biotin residues with high affinities, this likely reflects the high excess of streptavidin molecules, rapidly saturating available biotin residues. The distance between biotin residues may also be too large for significant crosslinking of the albumin network, reflecting the modest packing density of albumin molecules (packed monolayers are estimated between 190 and 630 ng/cm^2^, depending on the protein orientation).

**Figure 3.**
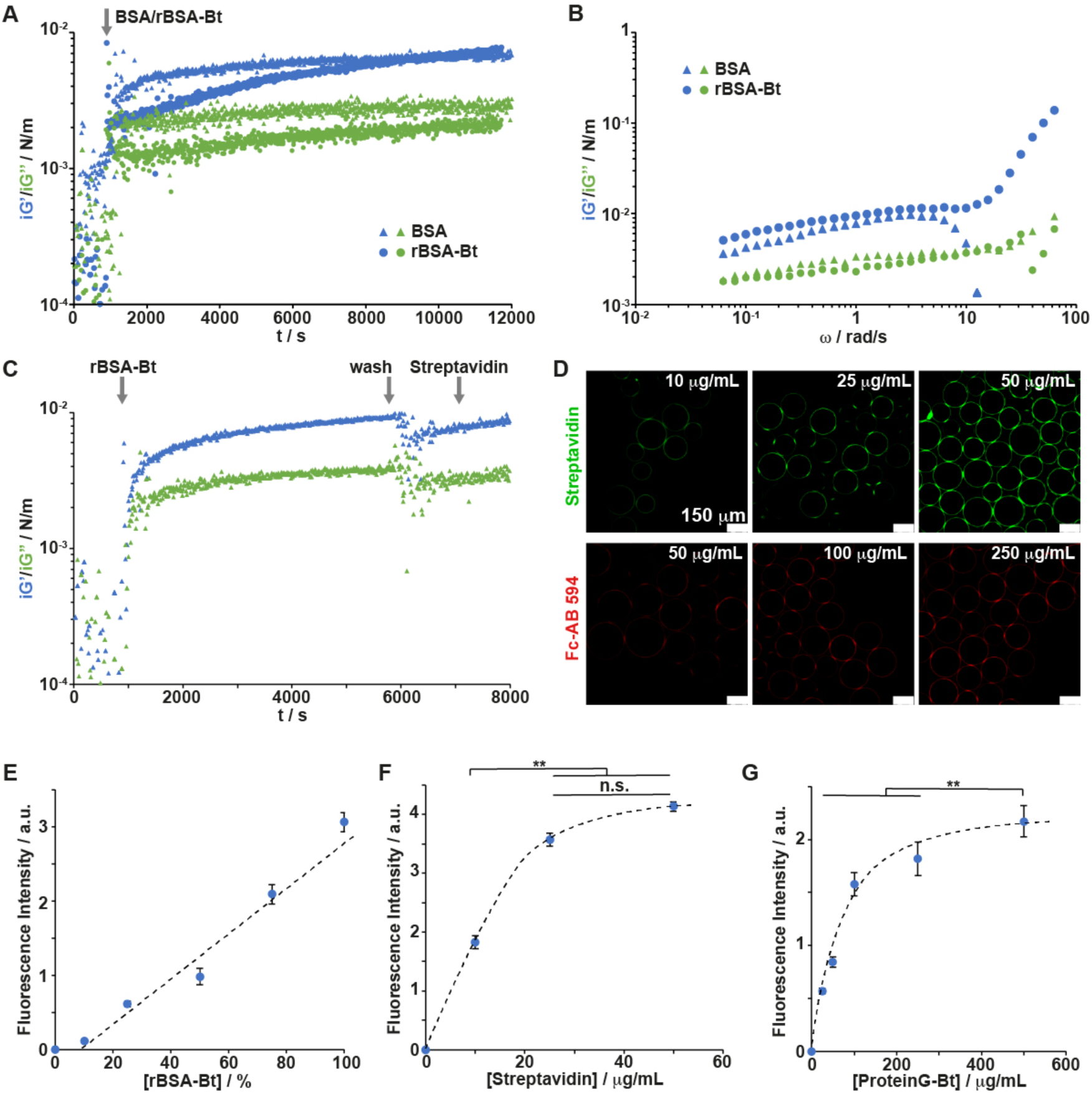
Formulation of artificial cells using engineered recombinant protein nanosheets. A. Representative time sweeps (frequency, 0.1 Hz; strain, 10^-3^ rad) monitoring the evolution of interfacial shear storage and loss moduli during the assembly of rBSA-Bt and BSA at the surface of Novec 7500 oil, in PBS (protein concentration: 1 mg/mL). The arrow indicates the time point of injection of corresponding proteins. B. Frequency sweeps of resulting interfaces (strain of 10^-3^ rad). C. Time sweep of sequential adsorption of rBSA-Bt, followed by streptavidin binding. The arrows indicate time points of injection of corresponding proteins. D. Fluorescence microscopy images of microdroplets of Novec 7500 stabilised by rBSA-Bt nanosheets, followed by streptavidin adsorption (green) at different concentrations and protein G (followed by 594-tagged antibody presenting Fc domains, shown in red, at a concentration of 4 μg/mL; protein G concentrations are indicated). E-G. The fluorescence intensities of streptavidin and tagged antibody ultimately detected (E and G) and streptavidin (F) were quantified as a function of starting concentration of rBSA-Bt (E), streptavidin (F) and Protein G-Bt (G). N = 3 ± SEM; n.s., not significant, p > 0.05; **, p < 0.01.

The formation of microdroplets by rBSA-Bt was next examined by microscopy (Figure 3D). Emulsions were formed by homogenisation of aqueous solutions containing albumins at a total concentration of 1 mg/mL with Novec 7500 oil. The mixing of rBSA-Bt with native BSA was investigated first, to identify whether biotin surface densities were a limiting factor for the subsequent binding of streptavidin. We observed a clear linear relationship between the concentration of rBSA-Bt and the ultimate antibody fluorescence intensity (pesenting Fc domains, Figure 3E), implying that the density of biotin residues, for emulsions formed in the presence of rBSA-Bt alone, was not saturated. This is in agreement with the SPR data (Figure 2D-E), which indicated higher molar surface densities of rBSA-Bt compared to streptavidin. Therefore, 1 mg/mL rBSA-Bt solutions were selected for further evaluation, to maximise streptavidin surface densities and the impact of the concentration of streptavidin was next examined.

Although the ultimate fluorescence intensity was found to increase significantly when the streptavidin bulk concentration was increased from 10 μg/mL to 25 μg/mL, it only increased modestly when the concentration was further increased to 50 μg/mL (Figure 3F). Therefore, an intermediate concentration was selected for further evaluation. Such low concentration, compared to the mg/mL concentration of albumins typically required for the stabilisation of emulsions is in agreement with the surface densities measured by SPR. Indeed, for 10 μm droplets, the weight of proteins required to stabilise the equivalent of 1 mL of oil phase is 0.44 mg. Therefore, at a 2/1 volume ratio of aqueous phase to oil, a modest excess of protein is used to stabilise emulsions. However, at a streptavidin density of 28 ng/cm^2^, compared to the 97 ng/cm^2^ measured for rBSA-Bt, the amount of streptavidin required to saturate the surface of 1 mL of 10 μm droplets is expected to be 126 μg. Therefore, in the conditions used for the functionalisation of artificial cells with the ligands of interest (10 μL of droplets for 100 μL of protein solution), a concentration of 25 μg/mL corresponds to a two-fold excess. Such efficiency is in good agreement with the very high binding affinity of streptavidin for biotin.

Finally, the impact of the concentration of biotinylated protein-G on the capture of Fc-tagged protein was characterised (Figure 3G). The intensity of antibody captured increased gradually as the concentration of ProteinG-Bt increased to 100 μg/mL and then reached a quasi-plateau at higher concentrations. This is again in good agreement with the surface density of ProteinG immobilised at the surface of nanosheets, as determined by SPR (61 ng/cm^2^). At such density, the mass of ProteinG required to saturate the surface of 1 mL of 10 μm droplets would be 280 μg and, at 100 μg/mL, with 10/1 volume ratio of aqueous to oil phases, a near four-fold excess was used in our experimental protocol. This larger excess required, compared to that observed for streptavidin adsorption, is to be expected considering the weaker binding affinity of subsequent biotin residues binding to streptavidin.

### 2.4. Embedding of artificial cells within embryoid bodies

The assembly of embryoid bodies with artificial cells was next examined. To do so, iPSCs were detached and resuspended with artificial cells functionalised with E-cadherin, N-cadherin or a mixture of both receptors, and allowed to adhere and form approximately 400-500 μm cellular assemblies. During this process, cells suspended at the bottom of a low adhesive plate were found to gradually make contact with other cells and cluster into a homogenous cell mass, with some artificial cells remaining entrapped (when presenting E-cadherins), whereas others were left at the bottom of the dish (Figure 4A). Incomplete uptake is to be expected from the higher density of the microdroplets formed, resulting in faster sedimentation and accumulation at the bottom of the culture dish. Following induction and differentiation, some of the droplets could be clearly seen at the periphery of developing organoids, whether at the surface or in the cell assembly (Figure 4B). The density of the core of the organoids did not allow resolving the deeper integration of artificial cells.

**Figure 4.**
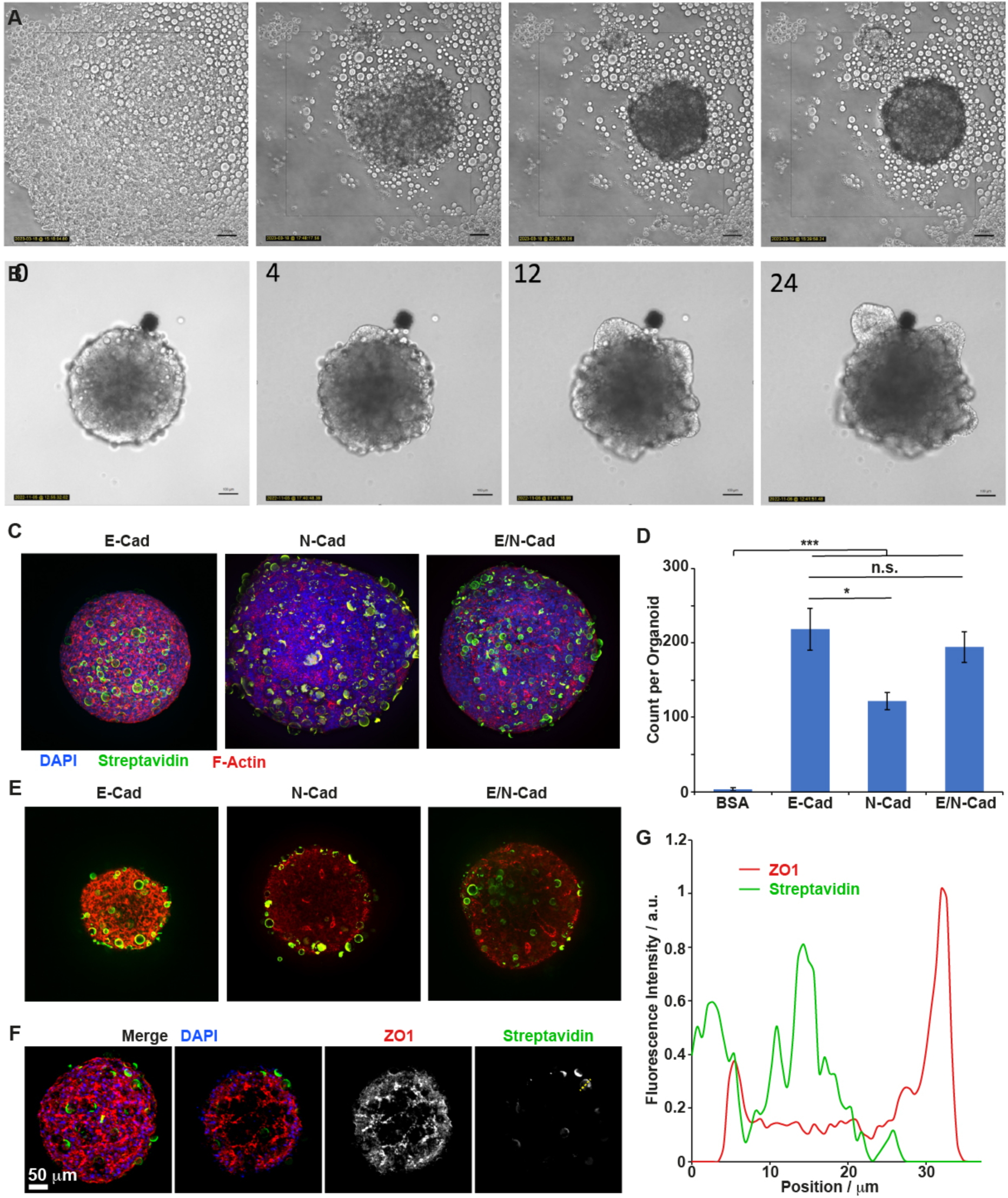
Integration of artificial cells within embryoid bodies. A. Series of frames taken from Supplementary Video S3, showing the integration of artificial cells (E-cadherin functionalised) by iPSCs during the formation of embryoid bodies. B. Series of frames taken from Supplementary Video S4, showing the morphology of developing organoids with integrated artificial cells (E-cadherin functionalised), following the induction and differentiation of embryoid bodies. C. Confocal microscopy images (Max projections of z-stacks) of embryoid bodies formed in the presence of artificial cells (Blue, DAPI; Green, Streptavidin; Red, F-actin). D. Corresponding quantification of counts of artificial cells per organoid. Native BSA was used as control. N > 5 ± SEM; n.s., not significant, p > 0.05; *, p < 0.05; ***, p < 0.001. E. Example of individual slices taken from confocal stacks of embryoid bodies formed in the presence of artificial cells (Green, Streptavidin; Red, ZO1). F. Example of stack and single slice imaging ZO1 localisation (Blue, DAPI; Green, Streptavidin; Red, ZO1), with separate channels shown. G. A profile of intensity (along the yellow dashed line shown in panel F) was measured across one of the artificial cells observed, showing no apparent localisation of ZO1 at its surface.

Confocal microscopy images, using tagged streptavidin to identify artificial cells, were used to evaluate their degree of integration within embryoid bodies (prior to induction, Figure 4C). The number of artificial cells per organoid was quantified for artificial cells presenting different types of cadherin receptors. In the absence of cadherins, no artificial cells could be observed. However, in the case of E-cadherin presenting artificial cells, high densities were detected (Figure 4D). In the case of N-cadherin-functionalised microdroplets, the densities were almost halved but recovered when mixed E- and N-cadherin presenting nanosheets were used to stabilise artificial cells. In addition, investigation of the separate sections revealed that, although relatively high densities of N-cadherin artificial cells were observed in the max projection of confocal stacks (to maximised 3D visualisation), most of these objects were found at the surface of the embryoid bodies (Figure 4E), with few found in the core. In contrast, E-cadherin presenting artificial cells were found both in the core and at the surface of cell assemblies. This observation is in good agreement with the expression of E-cadherin by iPSCs, whereas N-cadherin is not significantly expressed until ectodermal induction and neuroepithelial differentiation [58,82].

Finally, we examined confocal microscopy images of embryoid bodies prior to induction and the recruitment of the protein associated with the formation of tight junctions presenting ZO1 (Figure 4F). Although this protein was clearly recruited at junctions between cells forming the embryoid body, no clear recruitment was observed at the surface of E-cadherin presenting artificial cells, as was particularly apparent when inspecting intensity profiles of streptavidin and ZO1. Hence artificial cells were recruited within embryoid bodies and retained prior to ectodermal induction but did not induce the formation of mature cell-cell junctions. In addition, all artificial cells observed were spherical, with no evidence for significant deformation, implying that relatively little local stress was accumulating within embryoid bodies, which could have resulted in the distortion of microdroplets, as observed in the context of other cellular assemblies [26].

### 2.5. Impact of artificial cells on neuroepithelial organoid development

The fate of artificial cells during neuroepithelial differentiation was examined by induction of ectoderm commitment, followed by differentiation, as outlined in Figure 1A. At day 3 post induction, organoids displayed comparable numbers of neuroepithelial rosettes, in all conditions (Figure 5A-B). In addition, budding neuroepithelial rosettes displayed normal morphologies, with dimensions insensitive to the presence of artificial cells or the type of receptors they presented (Figure 5C). Further culture of organoids until day 9 led to the maturation of neuroepithelial organoids, with expression of SOX2 at neuroepithelia and initiation of the differentiation marker TUJ1 (Figure 5D). Therefore, neuroepithelial organoids initiated from artificial cell-loaded embryoid bodies were successfully induced and continued to mature normally, irrespective of the type of cadherin displayed, validating the proposed approach to introduce these objects within complex organoids, without disrupting their development.

**Figure 5.**
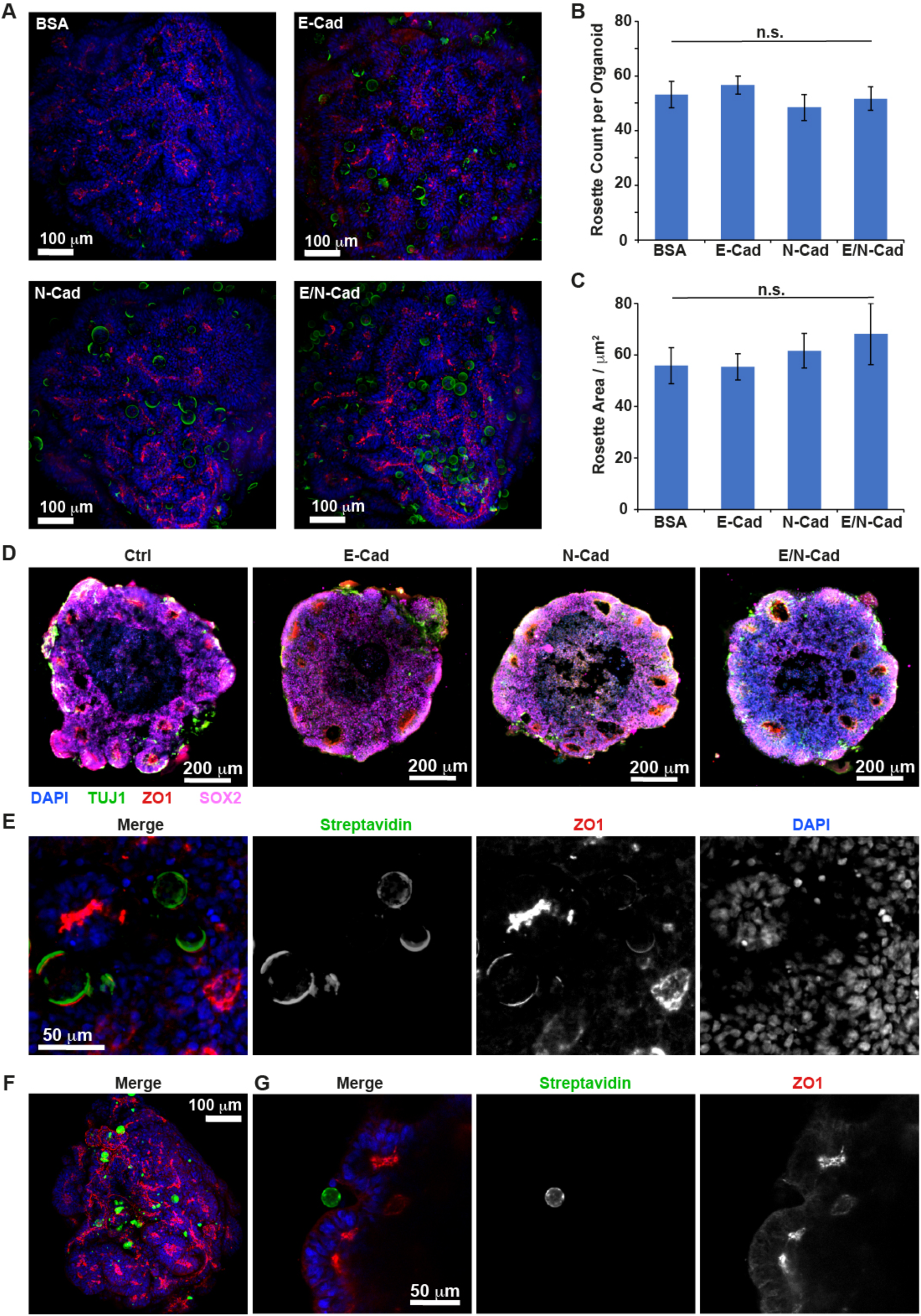
Rapid expulsion of large artificial cells from developing neuroepithelial organoids. A. Confocal microscopy images (Max projections of z-stacks) of developing neuroepithelial organoids (day 3) formed in the presence of artificial cells displaying different receptors (Blue, DAPI; Green, Streptavidin; Red, ZO1). B-C. Quantification of the number of rosettes (B) and their average area (C) observed per organoid. N > 6 ± SEM; n.s., not significant, p > 0.05. D. Confocal microscopy images (Max projections of z-stacks) of histological sections taken from developing neuroepithelial organoids (day 9) formed in the presence of artificial cells displaying different receptors (Blue, DAPI; Green, TUJ1; Red, ZO1; Magenta, SOX2). E. Example of individual optical slice taken from confocal stacks of developing neuroepithelial organoids (day 3) formed in the presence of E/N-cadherin presenting artificial cells (Blue, DAPI; Green, Streptavidin; Red, ZO1). F-G. (F) Confocal microscopy images (Max projections of z-stacks; left) and (G) example of zoom of individual optical slice taken from confocal stacks of developing neuroepithelial organoids (day 3) formed in the presence of silica microparticles (Blue, DAPI; Green, Streptavidin; Red, ZO1).

However, examination of the stacks did not reveal retention of artificial cells within neuroepithelia, and instead they accumulated in between rosettes, regardless of the type of receptor presented (Figure 5A). Further examination of the optical slices taken from confocal stacks confirmed that artificial cells were expelled from the budding neuroepithelial rosettes at day 3 (Figure 5E). Interestingly, at this time point, ZO1 was found to accumulate at the surface of artificial cells (Figure 5E). Although ZO1 localised at much higher densities at apical junctions in developing neuroepithelia, it was not recruited at lateral cell-cell junctions to the same extent to recruitment to artificial cells. Therefore, these data indicate that, upon induction of neuroepithelia formation, artificial cells are able to form junctions with surrounding cells, resulting in the accumulation of ZO1. However, despite the formation of junctions mimicking cell-cell interactions, artificial cells were rapidly expelled from developing neuroepithelia and localised in between budding rosettes. In addition, we could not see any evidence of droplet deformation, with all droplets observed appearing spherical and symmetrical (Figure 5E). Our data suggest that apical compressive forces arising within neuroepithelia rapidly expel artificial cells, resulting in their accumulation at their periphery.

To investigate the impact of artificial cell mechanics on their retention in neuroepithelia, we coated rigid silica microparticles (20 μm) with mixtures of E- and N-cadherins, using the same protocol used to stabilise artificial cells (silica particles were coated with hydrophobic octyl silane monolayers). Similarly to what was observed with artificial cells, the silica particles were rapidly expelled from the organoids, with few particles remaining adhered in between neuroepithelia (Figure 5F). This implies that rigid particles, as well as compliant droplets, are expelled from developing neuroepithelia, likely as a result of rapid proliferation in basal layers, and a build-up of apical compressive forces correlating with YAP cytoplasmic translocation. However, examination of ZO1 localisation in confocal slices revealed no association at the surface of microparticles, unlike what was observed with artificial cells (Figure 5G). This suggested that, although >20 μm cadherin-presenting objects were extruded from neuroepithelia regardless of their mechanical properties, the soft cell-cell adhesive interfaces of artificial cells may promote further maturation and biomimicry of cell-cell junctions.

### 2.6. Apical compressive forces expel artificial cell during neuroepithelial organoid early development

To further investigate the expulsion process evidenced with large artificial cells and microparticles (30 μm and 20 μm, respectively), we investigated the fate of 8 μm artificial cells and 5 μm silica particles. Indeed, mechanical forces exerted on smaller particles are expected to lead to slower expulsion. At day 3 post induction, a relatively large number of E/N-cadherin presenting artificial cells and silica particles remained within organoids (Figure 6A and Supplementary Figure S7). Compared to the densities measured initially within embryoid bodies, a significant reduction was still observed, indicating that, even for smaller objects, compressive forces resulted in the effective expulsion of a large proportion of artificial cells (Figure 6B). However, many particles and artificial cells were found remaining in rosettes, indicating a slower expulsion process. No significant differences were observed between the densities of artificial cells and those of silica nanoparticles remaining in organoids.

**Figure 6.**
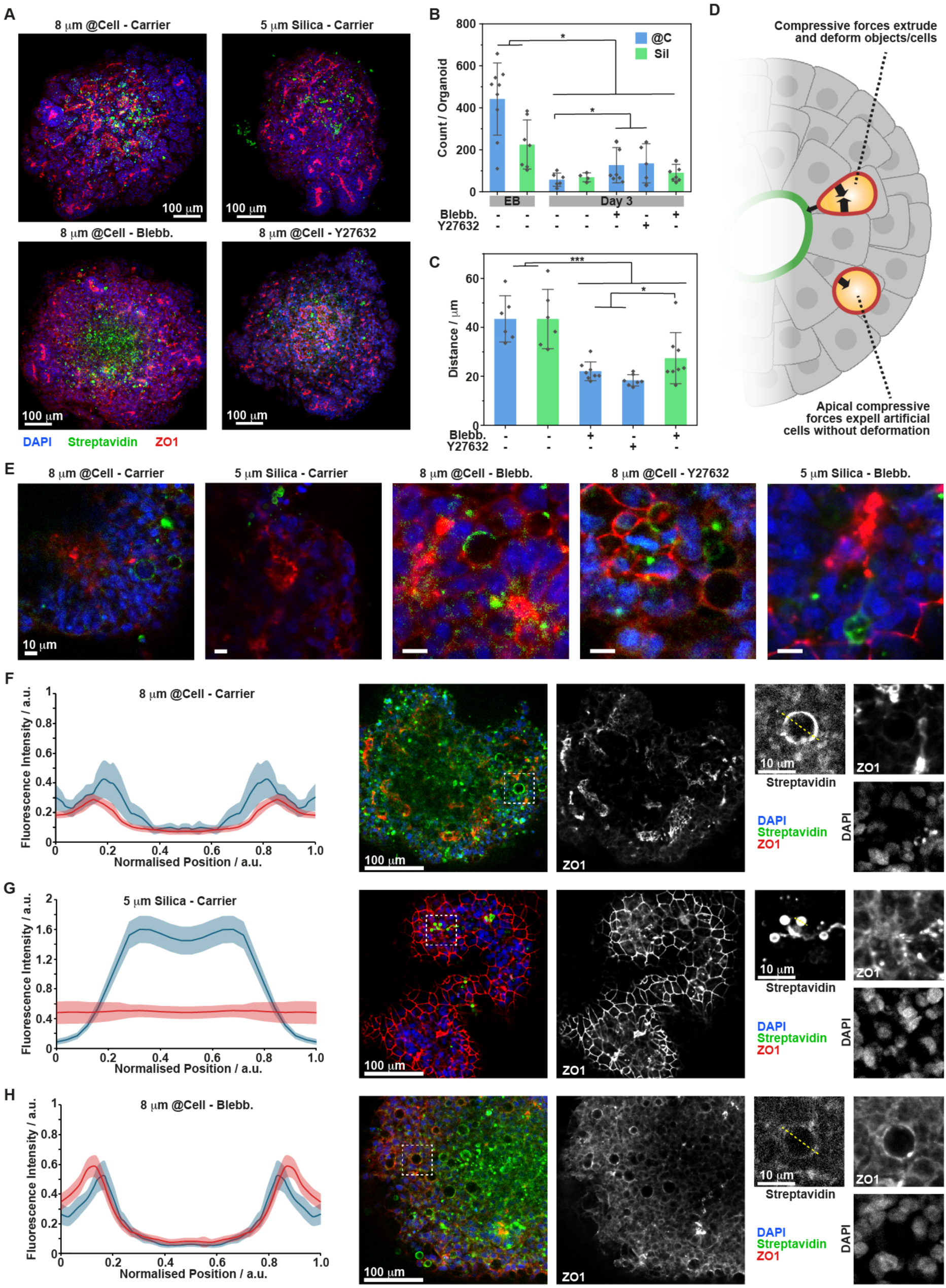
Artificial cells (8 μm) are retained by neuroepithelial organoids and promote ZO1 recruitment by neighbouring cells. A. Confocal microscopy images (Max projections of z-stacks) of neuroepithelial organoids seeded with 8 μm artificial cells (@Cell) or 5 μm silica microparticles presenting E/N-cadherins, at day 3 post induction, with and without Blebbistatin/Y27632 (10 and 20 μM, respectively; Blue, DAPI; Green, Streptavidin; Red, ZO1). Representative images obtained for other conditions are shown in Supplementary Figure S7. B-C. Corresponding quantification of counts of artificial cells/particles per organoid (B) and distance to the nearest ZO1+ cluster. N > 5 ± SEM; *, p < 0.05; ***, p < 0.001. D. Schematic representation of the proposed forces exerted on artificial cells, resulting in their expulsion. Two scenarios are proposed: in the first, shear forces/apical adhesion resist compressive forces and would result in artificial cell deformation and extrusion (not observed), whereas, in the second, apical compressive forces induce expulsion with negligible shear forces from basal cells (observed). E. Example of zooms taken from individual slices, from confocal stacks of day 3 organoids seeded with artificial cells or silica microparticles (see Supplementary Figure S8 for other representative images). F-H. Examples of single optical slices taken from stacks and zooms showing individual channels, together with the quantification of streptavidin (blue) and ZO1(red) intensity profiles (shaded areas correspond to associated standard deviations from >10 profiles). Organoids were seeded and cultured with the following: F, 8 μm artificial cells in the presence of carrier; G, 5 μm silica particles in the presence of carrier; H, 8 μm artificial cells in the presence of blebbistatin (10 μM).

To investigate the role of cytoskeleton assembly and contractility on the droplet/particle expulsion process, organoids were cultured in the presence of the ROCK inhibitor Y27632, and the myosin contractility inhibitor blebbistatin (Figure 6B-C and Supplementary Figure S7). These inhibitors were applied at the maximum concentration for which we observed early nucleation of ZO1 clusters underpinning early neuroepithelial rosette development (20 μM for Y27632 and 10 μM for blebbistatin). However, we note that for both inhibitors, beyond 3 days of culture, no further development of neuroepithelial rosettes could be observed. Therefore, these conditions only correspond to the perturbation of the very early onset of budding but are too disruptive to neuroepithelial development to continue further maturation.

In these conditions, we observed significantly higher artificial cells and silica particles being retained in day 3 organoids compared to controls (Figure 6C and E, and Supplementary Figure S8). In addition, we observed a significant number of artificial cells/particles being retained within neuroepithelial rosettes. This was quantified through the distance measured between the centre of the droplet/particle and the edge of the closest ZO1 positive cluster, corresponding to apical junctions of budding neuroepithelia (Figure 6C). In the presence of Y27632 or blebbistatin, the distance between artificial cells or microparticles and ZO1 clusters reduced significantly compared to untreated controls. A reduction in distance was also observed when comparing blebbistatin-treated organoids seeded with silica particles to those seeded with artificial cells. In these latter conditions, many artificial cells and silica microparticles were observed in direct contact with ZO1 clusters, or separated by 1 cell only, together indicating that the expulsion process had been considerably slowed down by disruption of actin assembly and myosin contractility. However, whether in control conditions or in the presence of inhibitors, the shape of artificial cells was spherical, with no evidence for anisotropic stress exerted at the surface of artificial cells. This contrasts with the significant deformation of droplets injected in cell aggregates and embryos [25,26]. Hence, our data suggest that internal actomyosin cytoskeleton-dependent stress building up within budding neuroepithelial organoids results in the expulsion of droplets and particles, with low resistance from peripheral basal cells/cell bodies, which would otherwise result in significant deformation of artificial cells, or as these are themselves expelled (Figure 6D). This also implies modest shear forces exerted by surrounding cells, so that droplet/particle expulsion occurs faster than the accumulation of local stress that could result in artificial cell deformation. Associated shear forces would be expected to increase significantly upon maturation of cell-cell junctions. Therefore, we further examined the developing organoids at day 3, focusing on the recruitment of ZO1.

We examined single optical sections taken from embryoid bodies (day 0) and organoids (day 3) and profiled the recruitment of ZO1 against the position of artificial cells or silica particles (Figure 6F-H and Supplementary Figure S9). Prior to induction, significant ZO1 recruitment was observed at the surface of artificial cells, often at higher densities to ZO1 distribution at neighbouring cell-cell contacts (Supplementary Figure S9). In comparison, relatively little ZO1 was recruited at the surface of silica microparticles. Upon induction of neuroepithelial development (at day 3), ZO1 localisation at the surface of artificial cells was significantly reduced, although still apparent, and remained basal for silica microparticles (Figure 6F-G and Supplementary Figure S9). A reduction of ZO1 recruitment and increase in apical compressive forces could account for the expulsion of droplets without significant distortion of their spherical shape.

Interestingly, in the presence of ROCK and myosin inhibitors, we observed significant recruitment of ZO1 at the surface of artificial cells at day 3 post-induction (Figure 6H and Supplementary Figure S9), compared to empty carrier. This implies that, although myosin-mediated contractility and F-actin cytoskeleton assembly are not critical to recruit ZO1 at cell-cell junctions (at the concentration selected), they can significantly strengthen ZO1 recruitment, resulting in its translocation to more contractile cellular junctions (although the direct measurement of cell-mediated contractile forces was not possible). In addition, the absence of significant ZO1 recruitment at the surface of silica microparticles (Figure 6G and Supplementary Figure S9) suggested that interfacial viscoelasticity (absent in the latter particles), and associated clustering or rearrangement of ligands, could play an important role in the strengthening of cell-cell adhesions and associated junction maturation.

## 3. Conclusions

The recreation of cellular and sub-cellular components enabling the capture of key cellular functions with synthetic building blocks offers striking opportunities for the understanding of living systems, but also the engineering of novel biotechnologies [83–86]. In this respect, extra-cellular matrix engineering has enabled exquisite control over a broad range of parameters, including integrin adhesions, growth factor stimulation, enzymatic degradation and mechanics [50,87]. Recently, a range of approaches have been proposed to allow recreating cell-cell interactions. However, introducing such cues as part of a matrix fails to capture the interfacial confinement, geometry and viscoelastic properties associated with ligands and receptors embedded in cell membranes. Artificial cells enabling to recreate such properties could allow to recreate some of the processes critical to the regulation of stem cell phenotype and the harnessing of their properties for regenerative medicine.

The present work demonstrates the formation of artificial cells able to trigger ZO1 recruitment in embryoid bodies and neuroepithelial organoids. The lack of recruitment to rigid silica particles highlights the importance of interfacial viscoelasticity in the regulation of cell-cell adhesion maturation. How viscoelasticity impacts on cadherin recruitment is unclear, but it is possible that such interfacial properties enable the clustering of ligands, whilst balancing tugging forces generated at developing junctions. Indeed, stress generated at cell-cell junctions are known to play a critical role on planar polarity in epithelia and tissue morphogenesis [6,49]. In turn, using artificial cells as force sensors, the present study provides evidence for apical compressive forces rapidly building up in the developing neuroepithelium, resulting in their expulsion to basal and peripheral compartments of organoids (Figure 7A). As the size of artificial cells increases, we propose that the speed at which they are expulsed increases. Similarly, perturbation of cytoskeleton assembly and contractility results in a reduction in compressive forces and a reduced rate of expulsion (Figure 7B). The lack of deformation of artificial cells suggests weak basal shear forces, resulting in expulsion without significant deformation. However, the role of surface tension and interfacial shear mechanics, which could mimic abnormally strong cortical mechanics (in comparison to interfacial mechanics in normal cell membranes), could also account, at least partly for the phenomena observed.

**Figure 7.**
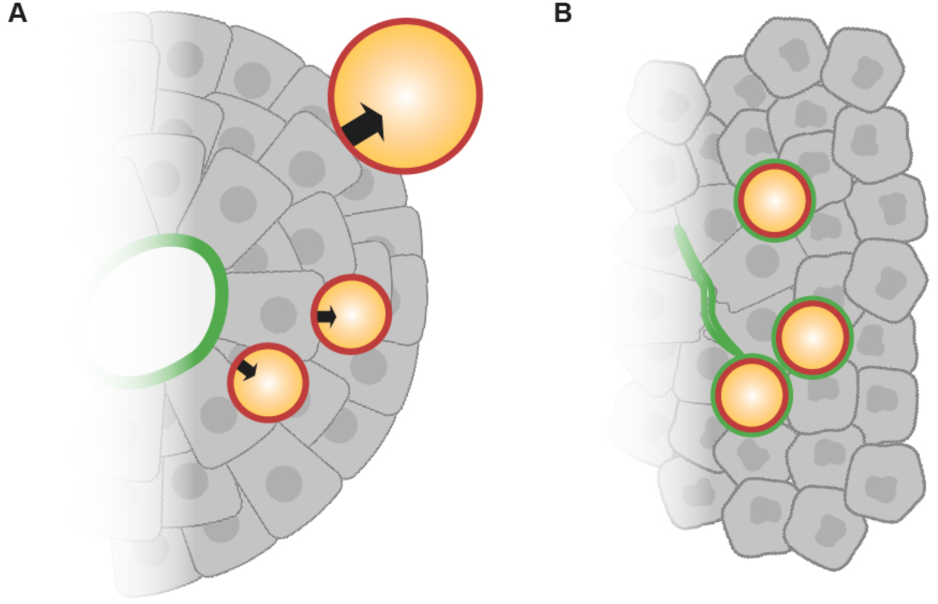
Summary of mechanical factors impacting on artificial cell expulsion from neuroepithelial organoids. A. Stronger apical compressive forces are exerted on larger artificial cells, resulting in faster expulsion. B. Disruption of cytoskeleton architecture and contractility results in impaired artificial cell expulsion, reflecting reduced compressive forces and increased cell-artificial cell junction maturation (indicated in green to reflect enhance ZO1 recruitment).

Overall, the joint observations that the translocation of the mechano-transducer YAP occurs rapidly upon induction of neuroepithelial organoid formation and that artificial cells and microparticles are rapidly expelled from developing neuroepithelia provides strong evidence for the occurrence of apical mechanical forces shaping the cerebral organoid early development. Although the origin of the tissue pressure and stresses that result in such morphological changes remains unclear, it is likely to be associated with the rapid cell expansion taking place, and cell divisions occurring at apical junctions [19]. As cell compression increases in the apical compartment, nuclei and cell bodies are gradually expelled outwards. In normal developing neuroepithelial organoids these forces are opposed by the retention of cell adhesion at apical junctions, leading to columnarisation and priming cell positioning for further differentiation and maturation. The combination of molecular biology and bioengineering tools, such as artificial cells, will allow shedding further light on these processes, in a human context.

## 4. Experimental Section

### 4.1. Materials

JM109 from Promega, Rosetta D3 (Novagen) from Sigma and DH5α *E. coli* cells from NEB. GS115 *P. pastoris* yeast strain from Life Technologies, Invitrogen. SHuffle® T7 Express Competent *E. coli* cells, Amylose resin and OneTaq® Quick-Load® 2X Master Mix were from New England Biolabs. The pPIC9K plasmid was from Gentaur Ltd. Restriction enzymes Fast Digest SnaBI, AvrII and SacI were from ThermoFisher. Ampicillin, DTT, glycerol, 100 mM ATP solution, D-biotin, 1 M MgCl_2_, PBS, PageRuler Unstained Broad Range Protein Ladder, Pierce biotin quantitation kit and Potassium phosphate dibasic were from ThermoFisher. D (+)-glucose, G-418, lithium acetate and PD10 desalting columns were from Sigma Aldrich. Agar, TAE 50X buffer and tris-glycine-SDS buffer, D-sorbitol, potassium phosphate monobasic and Lennox L Broth capsules, HEPES powder were from Melford Scientific. Casamino acid powder, Yeast extract, peptone and Yeast synthetic drop out medium (without histidine) was ordered from ForMedium. 5M NaCl was from Invitrogen. Imidazole powder was Acros Organics.

### 4.2. Plasmid design and cloning

The BSA protein sequence used for engineering rBSA-Avi was the 3V03 *Bos Taurus* sequence. The expression vector 6xHis-3C-Avi-rBSA was designed using the SnapGene software and ordered via the ThermoFisher GeneArt services. The *Bos Taurus* BSA codons were optimised using this service to the *P. pastoris* host codons to ensure reliability in expression. The inserts were then ordered and synthesised from ThermoFisher in a standard pEX-A2 vector containing the Amp® ampicillin resistance gene. 1 µL of pEX-A2-rBSA-Avi vector was transformed into 50 µL of JM109 bacteria via standard heat shock at 42°C and plated onto 100 µg/mL ampicillin LB plates overnight at 37°C. One colony was picked, transferred into 5 mL of 100 µg/mL ampicillin LB media and incubated overnight at 37°C, 220 rpm. 2 mL of the resulting starter culture was centrifuged at 14k rpm and the pellet was mini-prepped using a QIAGEN QIAprep miniprep kit with a final elution volume of 50 µL. pEX-A2-BSA-Avi and pPIC9K vectors were both digested with SnaBI and AvrII restriction enzymes at 37°C for 15 min using the digestion master mix in Supplementary Table S1. 20 µL of the digested plasmids were characterised on a 1% agarose gel for 30 minutes at 120V. 5 µL of DNA ladder were also used and loaded into the first lane. The recombinant BSA (rBSA) insert (1852 bp) and pPIC9K (9276 bp) were purified out of 1% agarose gel using a GeneJET gel extraction kit with a final elution volume of 50 µL. 4.5 µL of digested rBSA-Avi was mixed with 0.5 µL of digested pPIC9K and 5 µL of Instant sticky end master mix. 5 µL of the resultant ligated pPIC9K-rBSA-Avi vector was transformed into DH5α *E. coli* (high competency) cells using the standard heat shock procedure and 200 µL of the transformation was plated onto a 100 µg/mL ampicillin LB plate and incubated overnight at 37°C. Six colonies were picked from the plates to form 5 mL starter cultures and subsequently 2 mL of culture was miniprepped. 500 µL of each starter culture was mixed with 500 µL of 30% glycerol, vortexed, snap frozen with liquid nitrogen then stored at -80°C. Analytical digests were incubated at 37°C for 15 min using the digestion mix in Supplementary Table S2, and were setup for the miniprepped plasmids, before characterisation on 1% agarose gels at 120V for 30 min. Successful transformation was confirmed on agarose gels (Supplementary Figure S3), as two bands corresponding to the cleaved empty pPIC9K plasmid (9.27 kbp) and the rBSA-Avi insert (1.85 kbp). 10 µL of successful plasmids were sent off for sequencing to further verify the sequence along with 10 µM 5’ AOX1 forward primer. A successful transformant was streaked onto a 100 µg/mL ampicillin LB plate and incubated at 37°C overnight. Colonies were picked and transferred into 50 mL 100 µg/mL ampicillin LB media and incubated overnight at 37°C. The culture was midi-prepped using the QIAGEN plasmid plus midi prep kit to obtain at least 10 µg of plasmid DNA. Plasmids were stored at - 20°C until required.

### 4.3. Generating electrocompetent P. pastoris cells

GS115 *Pichia pastoris* strain was streaked onto a YPD-agar plate and incubated at 30°C for 2 days. Single colonies were then streaked into patches of roughly 4 cm^2^ onto YPD-agar and incubated at 30°C for 18 h. 920 µL of YPD was prepared in a sterile 1.5 mL Eppendorf tube, 40 µL of fresh 1M DTT and 40 µL of 1M HEPES-NaOH (pH 8.0) was added to the mix. Cells from one patch were scraped and transferred to the mix using a sterile pipette tip, the mix was vortexed to allow for uniform suspension. The tube was incubated at 30°C for 15 min at 100 rpm, then pelleted at 3000 x g for 3 min. The cells were washed twice with ice cold sterile water then twice with ice cold 1M sorbitol, with a final volume of 50 µL. The *P. pastoris* cells are now considered electrocompetent and can be stored at -80°C for long term storage.

### 4.4. Transformation of pPIC9K into P. pastoris

The pPIC9K-rBSA-Avi plasmid was digested and linearised by SacI restriction enzyme using the conditions in Supplementary Table S3. The linearised plasmid was then purified using the GeneJET PCR purification kit. The electrocompetent cells, along with 1 μg of linearised pPIC9K-rBSA-Avi were transferred to a pre-chilled electroporation cuvette and incubated on ice for 5 min. The moisture was wiped off the cuvette and pulsed at 2000 V/200 Ω/25 μF, followed by the immediate addition of 1 mL of ice cold 1 M sorbitol to the cuvette for recovery. The cells were transferred to a sterile 1.5 mL Eppendorf tube and incubated at 30°C for 1 h with no shaking. After 1 h, 500 μL of YPD was added to 500 μL of recovered cells and incubated overnight at room temperature to acquire G418 antibiotic resistance. The cells were pelleted at 3000 x g for 3 min and resuspended in 250 μL of water. 50 μL of cells was plated onto increasing concentration of G418 YPD-agar plates 0.25, 0.5, 1.0, 2.0, 4.0 mg/mL G418 and incubated at 30°C for 4-5 d dependent on antibiotic concentrations. Colonies from the highest antibiotic concentration were transferred to 5 mL YPD medium and starter cultures were generated overnight at 30°C, of which 500 μL was used to generate glycerol (15%) stocks. 10 μL of each starter culture was used to streak onto YNBD plates for His^+^ selection to eliminate any spontaneous G418 colonies.

### 4.5. Colony PCR of successful transformants

Successful colonies that showed both the highest antibiotic resistance and His+ phenotype were used for colony PCR (primers in Supplementary Table S4), this was conducted to check for the insertion of the pPIC9K-rBSA-Avi cassette at the chromosomal AOX1 promoter. In a 1.5 mL Eppendorf tube, 200 μL of liquid starter culture was spun down at 14,000 rpm for 1 min. The pellet was resuspended in 100 μL of 200 mM LiOAc + 1% SDS buffer and incubated at 70°C for 5 min. 300 μL of 100% ethanol was added and vortexed for 10 s. The DNA was spun down at 13,000 rpm for 3 min and the supernatant was discarded carefully without disrupting the white pellet. The pellet was washed once with 500 μL of 70% ethanol and spun down at 13,000 rpm for 1 min. The ethanol was removed without disrupting the pellet and dissolved in 100 μL RNA free ddH2O, then spun at 13,000 rpm for 1 min. 2 μL of the supernatant was used for PCR using the PCR mix in Supplementary Table S5. In the case of a Mut^+^ His^+^ phenotype, two bands should appear once the PCR product is run on a 0.8% agarose gel at 120V for 30 min. One band corresponding to the chromosomal AOX1 promoter at 2.2 kb and another band corresponding to the rBSA-Avi insert sequence plus the 492 bp PCR fragment generated by the parent plasmid at 2.34 kb. Successful transformants that observed both bands were used for subsequent protein expression.

### 4.6. Expression of rBSA-Avi in P. pastoris

A successful transformant from the highest concentration of antibiotics was streaked onto YPD-Agar containing 4.0 mg/mL G418 and incubated at 30°C for 3 d. One colony was transferred per 25 mL of BMGY medium in a falcon tube and incubated at 30°C, 220 rpm overnight. The cells were pelleted at 1500 x g for 5 min, resuspended in 5 mL of BMMY medium and transferred into a 2 L flask containing 500 mL of BMMY medium. The flasks were covered in a layer of sterile gauze and incubated at 28°C, 200 rpm for four days; 2% methanol was supplemented into the medium every 24 h to maintain induction of the AOX1 promoter and protein expression. A 1 mL sample was taken every 24 h to be used for the SDS-page gel. After four days, the cells were centrifuged at 7000 rpm for 15 min and the supernatant was further centrifuged in batches at 15,000 rpm for 2 min. The supernatant was then filtered and stored at 4°C until purification. Two 5 mL HisTrap columns used in series were connected to an AKTA Pure system and equilibrated using 5 column volumes of wash buffer (PBS + 20 mM imidazole). The supernatant containing the expression rBSA protein was loaded and applied to the column, then washed using 20 column volumes of wash buffer. The protein was eluted from the column using elution buffer (PBS + 500 mM imidazole), the elution fractions were pooled and stored at 4°C. The column was regenerated with three column volumes of ddH_2_O, followed by three column volumes of 1 M NaCl + 0.1 M NaOH, five column volumes of ddH_2_O and finally stored in 20% EtOH. SDS samples were taken at every step and a SDS gel was run at 220V for 30 minutes to verify protein purification. The excess salts and imidazole were removed from the buffer using a dialysis step through 14K MWCO dialysis tubing. 5 L of PBS was prepared in a container and the protein solution was pipetted into the dialysis tubing, tied from both ends and allowed to dialyse overnight. The rBSA-Avi protein was stored at -80°C in 5% glycerol for long term storage or used immediately for biotinylation.

### 4.7. Biotinylation of rBSA-Avi

If the Avi-tagged rBSA protein was thawed from long term storage, the 5% glycerol was removed prior to biotinylation using a PD10 desalting column and eluted using 3.5 mL of PBS buffer. The protein was further concentrated using a 30k MWCO concentrator to reach the 40 μM threshold required for optimal biotinylation. The O/D_280_ was measured for Avi-rBSA using a Quartz cuvette at 280nm in PBS buffer, and the protein molarity was calculated. Biotinylation was setup in a sterile 15 mL Falcon tube using filtered reagents according to the conditions stated in Supplementary Table S6. The biotinylation mix was incubated at 30°C, 100 rpm for 1 h. Amylose resin was washed with PBS three times by centrifugation and 100 µL of 50% amylose resin slurry was added to the mix and rocked for 30 min at room temperature. The mix was centrifuged at 14,000 rpm for 2 min and the supernatant was transferred to a 15 mL Falcon tube. 2 mL of sample was run once more through a PD10 desalting column to remove excess salts and biotin and eluted into 3.5 mL PBS. The sample was concentrated once more using a 30k MWCO and the protein molarity was measured using a Quartz cuvette at 280 nm in PBS. Three methods were used to measure the extent of biotinylation. As a quantitative method, the Pierce Biotin Quantitation Kit was used to quantify the amount of biotin molecules per molecule of protein; with a bt-HRP protein used as control. As a qualitative method, 5 µL of 10 µM rBSA-Bt was added to 10 µL of 2x SDS buffer, the sample was heated at 95°C for 5 min and allowed to cool to room temperature. 5 µL of 1 mg/mL streptavidin was added and incubated for 10 min at room temperature. The sample was run on an SDS-page gel at 220V for 30 min and the bands were observed for an increase in molecular weight. As a secondary qualitative method, emulsions were generated with the rBSA-Bt, washed with buffer and then incubated with 50 µg/mL FITC-Streptavidin and imaged (at 488 nm) to verify streptavidin binding.

### 4.8. Circular dichroism (CD) and mass spectrometry

rBSA-Bt protein was concentrated to 150 µM using a 30 kDa MWCO PES concentrator (Vivaspin) run at 4000xg and subsequently 1 mL of protein was loaded onto a S200 10/300 SEC column (Cytvia), preequilibrated with 5 mM potassium phosphate buffer pH 7.4, the flow rate was set to 0.25 mL/min and run overnight. This step was completed to exchange the buffer from PBS into IPO_4_ – a buffer that will not strongly absorb in the UV wavelength range of 180-300 nm. The CD instrument was purged with nitrogen gas. The temperature was set to 20°C. The photomultiplier voltage was measured at 300 nm with a sample chamber containing IPO_4_ buffer. 300 µL of 5 µM protein was transferred into a cuvette and loaded into the CD instrument (with a pathlength of 0.1 cm) and a wavelength scan protocol was conducted in the 180-300 nm range. Mass spectroscopy was carried out using the JEOL JMS-S3000 SpiralTOF™ MALDI-TOF mass spectrometer, mass spectra were acquired in SpiralTOF mode with a mass range of 23,000–82,000 m/z.

### 4.9. Surface plasmon resonance

All SPR measurements were carried out on a BIACORE X from Biacore AB. Gold coated SPR chips measuring 10 x 12 mm in size were plasma oxidised for five minutes using the Henniker Plasma HPT-200 machine. Then treated in a 5% methanolic solution of 1H,1H,2H,2H-perfluorodecanethiol overnight at room temperature. This generated a fluorinated monolayer which served as a model to mimic the fluorinated Novec-7500 oil. The chips were then washed once with water and dried in air prior to mounting. The sample sensor chip was then mounted onto a plastic support frame, docked and primed once with PBS. Once primed the signal was allowed to stabilise to a stable baseline. rBSA-Bt (1 mg/mL) was then loaded into the sample loop with a micropipette and injected at a flow rate of 10 μL/min. Once injection had finished, the surface of the sample chip was washed with PBS for 10 min at a flow rate of 10 μL/min followed by the subsequent injection of streptavidin (50 μg/mL). The injection process was repeated with biotinylated protein G (100 μg/mL), followed by Fc-E-cadherin (25 μg/mL).

### 4.9. Interfacial shear rheology

Interfacial shear rheological measurements were carried out on a Discovery Hybrid Rheometer (DHR-3) from TA Instruments. The diamond-shaped Du Noüy Ring (DDR) geometry used had a radius of 10 mm and was made of platinum-iridium, with a diameter of 0.36 mm. 4 mL of Novec-7500 fluorinated oil was pipetted into the circular double walled trough. Using Axial force monitoring, the ring was positioned at the surface of the fluorinated oil interface, a further 180 μm was subtracted from the contact point to position the medial plane of the ring at the interface. The trough was then filled with 4 mL of PBS solution to fully cover the oil subphase. A time sweep with a constant frequency of 0.1 Hz, a temperature of 25°C and a displacement of 1.0 10^-3^ rad was run for a total of 12,000 s. After 900 s, a solution of rBSA-Bt was injected into the aqueous phase at a final concentration of 1 mg/mL and left to self-assemble at the interface for 1 h. After 1 h, a frequency sweep with a displacement of 1.0 10^-3^ rad was carried out. The measurement was paused and the aqueous phase was then washed with 30 mL of PBS using an Elveflow system at 1 mL/min flowrate for 30 min. The interface was given a further 5 min to stabilise and a solution of streptavidin was injected into the aqueous phase at a final concentration of 100 µg/mL. Time and frequency sweeps were repeated.

### 4.10. iPSC culture

5700 μL of DMEM/F-12 was mixed with 300 μL of thawed Corning® Matrigel® (GFR) Basement Membrane Matrix. 1000 μL of the mix was then pipetted into three wells of two separate 6 well Falcon plates, for a total of six coated wells, swirled to ensure full coverage and incubated at 37°C for 1 h. After 1 h, one plate was used for experiments and the other stored at 4°C; coated plates should not be stored for more than a week. Human iPSCs (hIPSCs) were cultured in standard incubator conditions of 5% CO_2_ at 37°C. After 24 h post-thawing, the rock-E8 media was removed and replaced with 4 mL of complete E8 Flex medium (no rock inhibitor). The media was exchanged every 2 days until cultures reached approximately 80% confluency. Areas of spontaneous differentiation or embryoid body formation were removed before the passaging process – to do this the media was removed and 1 mL of dPBS was gently pipetted directly onto the affected area to detach the differentiated cells (using a microscope to view colonies). Cell detachment solution for passaging was prepared in a 1.5 mL Eppendorf tube by mixing 1 μL of 0.5M EDTA (pH 8.0) with 1 mL of dPBS. Media were removed from the well and replaced with 1 mL of cell detachment solution and incubated at 37°C for 3.5 min. The solution was slowly pipetted up and down 3-4 times using a 5 mL stripette onto the cells to break up the culture. The cells were then transferred into a sterile 15 mL Falcon tube and centrifuged at 300 x g for 2 min. A fresh Matrigel plate was prepared by removing the existing DMEM/F12-Matrigel solution, washing once with dPBS and replacing with 3 mL of complete E8 Flex medium. The supernatant was removed carefully using a 1 mL pipette to not disrupt the cell pellet and resuspended into 3 mL of complete E8 Flex medium (Supplementary Table S7 Gibco™). The cell suspension was evenly distributed between the three wells and incubated at 37°C for 3-4 days; the cells were observed under the microscope to verify that clusters of cells were present before incubation. The medium was exchanged every 2 d until 80-90% confluency was obtained, then the passaging cycle was repeated. hIPSCs were used until a cell passage number of 13 or until cells began to proliferate at a slower rate (taking more than 4-5 d to reach 80% confluency).

### 4.11. Formation of embryoid bodies

Once the hIPSCs maintenance culture reached a confluency of 80%, the media was removed from one well and washed once with dPBS. 1 mL of Gentle dissociation reagent (Stem cell technologies) (Supplementary Table S8) was added into the well and incubated at 37°C for 7.5 minutes. The cells were broken gently using a 5ml stripette at low setting 4-5 times and transferred to a sterile 15 mL Falcon tube. 2 mL of embryoid body seeding medium was added to the cells and centrifuged at 300 x g for 2 minutes. The supernatant was removed carefully not disrupting the pellet and resuspended in 1 mL of EB seeding medium. 10 μL was transferred to a haemocytometer and the average cell number/mL was quantified. The cells were resuspended into 3 mL of embryoid body seeding medium with a final cell density of 90,000 cells/mL. 100μL of cell suspension was added into each well of a sterile 96 well ultra-low attachment (ULA) plate – final cell density per well of 9000 cells. The plate was spun at 700 rpm for 30 s, at room temperature, to pull the cells into the centre of each well. The plate was incubated at 37°C without disruption for the first 24 h. 100 μL of embryoid body formation medium was pipetted gently into each well and incubated for 2 days.

### 4.12. Generation of neuroepithelial organoids

At day 0, embryoid bodies were induced to neuro-ectodermal lineages. Using a 1000 μL pipette tip, the embryoid body media were carefully removed from each well, leaving behind the embryoid body in the well (roughly 4-5 wells per tip). 200 μL of induction medium (Supplementary Table S9) was then pipetted onto the walls of each well with minimal disruption of the embryoid body. The plate was incubated for another 2 d at 37°C. At day 2, the differentiation medium was prepared and the thawed Matrigel was added freshly, immediately before feeding. Using a 1000 μL pipette tip, the induction media was removed from each well, leaving behind the organoid in the well (roughly 4-5 wells per tip). 200 μL of differentiation medium was then pipetted into each well with minimal disruption of the EB. The plate was further incubated for 3 d. At day 3, the EB maturation medium was prepared and the thawed Matrigel was added freshly, immediately before feeding. Using a 1000 μL pipette tip, the expansion media was removed from each well, leaving behind the organoid in the well (roughly 4-5 wells per tip). 200 μL of organoid maturation medium was then pipetted into each well with minimal disruption of the organoid. The plate was further incubated for 6 days. To investigate the impact of ROCK and myosin inhibition, the ROCK inhibitor Y27632 and blebbistatin were used, respectively, at concentrations of 20 μM and 10 μM (following evaluation of a range of concentrations of 4-20 μM and 10-50 μM, respectively.

### 4.13. Formation of artificial cells and coating of silica microparticles

2 mL of filtered 1 mg/mL rBSA-Bt solution was added to 1 mL of filtered Novec-7500 oil inside a glass container. The head of a T 25 digital ULTRA-TURRAX homogenizer-emulsifier dispersion mixer from IKA was lowered into the solution and spun at varying speeds depending on droplet sizes (15,000 rpm for 30 μm and 20,000 rpm for 8 μm), for 2 min. The resulting emulsions were left to recover for 5 min, then transferred using a 3% BSA coated pipette tip into a 3% BSA coated 15 mL Falcon tube. 100 µL of emulsions were transferred to a 3% BSA coated sterile 1.5 mL Eppendorf tube and washed six times with PBS. The final concentrations used for functionalisation for seeding into organoids were the following: 50 µg/mL FITC-Avidin (Thermo Fisher, 21221), 100 µg/mL Bt-ProteinG (Thermo Fisher, 29988), 25 µg/mL Fc-tagged E-Cadherin (R&D Systems, 648-EC) and 25 µg/mL Fc-tagged N-Cadherin (R&D Systems, 1388-NC). Excess PBS was removed from the emulsions and 100 µL of 50 µg/mL of FITC-Streptavidin was added and incubated at room temperature for 15 min. The emulsions were washed six times with PBS and imaged via fluorescence microscopy (at 488 nm) to verify fluorescence. After washing with PBS twice, the PBS was exchanged with 100 µL of 100 µg/mL bt-ProteinG and incubated at RT for 15 min. The emulsions were washed six times with PBS and the PBS was exchanged with a sterile 25 µg/mL Fc-tagged N-cadherin or E-Cadherin solution (or a 1/1 mixture of the two), then incubated for 1 hour at 4°C. The emulsions were washed six times with PBS and 1 mL of dPBS was added to a 15 mL Eppendorf tube. For Silica particle functionalisation, silica particles (coated with octyl silane, Sigma) with a diameter of 5 and 20 µm were functionalised using the same protocol as the oil droplets, using 50 µg/mL FITC-Streptavidin, 100 µg/mL Bt-ProteinG, 25 µg/mL Fc-tagged E-Cadherin and 25 µg/mL Fc-tagged N-Cadherin.

### 4.14. Formation of embryoid bodies in the presence of artificial cells or particles

Artificial cells and silica microparticles were mixed with single cell hIPSCs at day -2 to form embryoid bodies. Once the hIPSCs were seeded at a density of 4500 cells/well and centrifuged to create a cell mass in the centre of the well, a 1000 µL pipette tip coated with 3% BSA was used to partially resuspend the emulsions in the dPBS and using a 10 µL pipette tip coated with 3% BSA, 1 µL of emulsions (or silica particles) was transferred directly on top of the cell masses in each well. The embryoid bodies and organoids were cultured as per the protocols reported above.

### 4.15. Cryosectioning of organoids

At day 9 of organoid culture, the organoids were fixed using 4% PFA for 15 min and washed six times with PBS. The organoids were placed within the central region of a mould containing Optimal cutting temperature (OCT) compound, and snap frozen using a hexadecane ice bath. The blocks were stored at -80°C, until required. The blocks were cryosectioned using an OTF7000 Cryostat with Epredia™ Microtome Blades at -10°C ambient temperature and -20°C sample temperature, using a slice thickness of 25 µm. Slices were placed onto frost microscope slides and stored at -20°C until required.

### 4.16. Immunostaining and imaging

Embryoid bodies and organoids were fixed using 4% PFA treatment for 20 min at room temperature, followed by washing with PBS and treatment with 0.25% Triton-x for 15 min at room temperature, followed by PBS washing. The organoids were washed six times with PBS and blocked with 3% BSA for 30 min and incubated with primary antibodies for 1 h at 4°C. The organoids were washed six times with PBS and incubated with secondary antibodies, Phalloidin-555 and DAPI for 1 h at RT. The organoids were washed six times with PBS and transferred using a 3% BSA coated 1000 µL pipette tip into a PBS filled Ibidi µ-Slide 8 Well Chamber coated in 3% BSA. For organoids cultured for 9 days and cryosectioned, slides containing cryosectioned organoid slices were thawed from -80°C. Using a hydrophobic marker, a square was marked multiple times around the slices. 50 µL of 4% PFA was pipetted into the seal directly on top of the slices and incubated for 15 minutes at room temperature. The slices were washed three times with PBS and exchanged with 50 µL of 0.25% Triton-X and incubated for 15 min at room temperature. The slices were washed three times with PBS and blocked with 3% BSA for 30 min and incubated with primary antibodies for 1 h at 4°C. The slices were washed three times with PBS and incubated with secondary antibodies, Phalloidin-555 (Thermo Fisher, R415) and DAPI-408 for 1 h, at room temperature. The slices were washed three times with PBS and the PBS was replaced with a small drop of Fluoromount-G™ Mounting Medium. A circular thin glass slip was placed on top of each slice and left to set for 30 min. Details of antibodies and staining reagents: anti-Tubulin β3 (BioLegend, Alexa Fluor® 488, 801203); YAP1 (C-terminal, Sigma, Y4770); rhodamine phalloidin (Invitrogen, R415); DAPI (Invitrogen, D1306); SOX2 (Invitrogen, Polyclonal Antibody, PA1-094); ZO-1 (ThermoFisher, Alexa Fluor™ 488, ZO1-1A12), PAX6 (ThermoFisher, Alexa Fluor™ 594, 42-6600); FOXG1 (Abcam, ab18259); secondary antibodies were from Invitrogen (A-31573, A-31571, A-11008, A21436, A31571, A11012, A-11029, A31570, A-31573, A21203). Slides were imaged on a Nikon CSU-W1 SoRa Spinning Disk Confocal microscope with a pinhole of 50 µm and controlled via NIS-Elements software, a 20X air objective was used for the majority of imaging. Samples were illuminated using lasers at 405 nm (DAPI), 488 nm (GFP), and 561 nm (mCherry) with laser powers set to a range of 10% and exposure times of 500 ms per channel.

## Supporting Information

Supporting Information is available from the author.

## Data Availability

All data analysed during this study are included in this published article (and its supplementary information file). Other raw data required to reproduce these findings are available from the corresponding author on reasonable request.

## Acknowledgements

We thank Dr Madeline Lancaster for insightful discussions on the development or cerebral organoids and their derivation, and Ms Arushi Metha for assistance with imaging of organoids. Funding for this work from the European Research Council (ProLiCell, 772462, ProBioFac, 966740 and ProNaGen, 101138464) is gratefully acknowledged.

## References

[1] D.P. Kiehart, J.M. Crawford, A. Aristotelous, S. Venakides, G.S. Edwards, Cell sheet morphogenesis: Dorsal closure in drosophila melanogaster as a model system, Annu Rev Cell Dev Biol 33 (2017) 169–202. 10.1146/annurev-cellbio-111315-125357.

[2] C.J. Miller, L.A. Davidson, The interplay between cell signalling and mechanics in developmental processes, Nat Rev Genet 14 (2013) 733–744. 10.1038/nrg3513.

[3] R. Keller, L.A. Davidson, D.R. Shook, How we are shaped: The biomechanics of gastrulation, Differentiation 71 (2003) 171–205. 10.1046/j.1432-0436.2003.710301.x.

[4] A.C. Martin, M. Gelbart, R. Fernandez-Gonzalez, M. Kaschube, E.F. Wieschaus, Integration of contractile forces during tissue invagination, Journal of Cell Biology 188 (2010) 735–749. 10.1083/jcb.200910099.

[5] D.S. Vijayraghavan, L.A. Davidson, Mechanics of neurulation: From classical to current perspectives on the physical mechanics that shape, fold, and form the neural tube, Birth Defects Res 109 (2017) 153–168. 10.1002/bdra.23557.

[6] A.C. Martin, B. Goldstein, Apical constriction: Themes and variations on a cellular mechanism driving morphogenesis, Development (Cambridge) 141 (2014) 1987–1998. 10.1242/dev.102228.

[7] Y. Inoue, M. Suzuki, T. Watanabe, N. Yasue, I. Tateo, T. Adachi, N. Ueno, Mechanical roles of apical constriction, cell elongation, and cell migration during neural tube formation in Xenopus, Biomech Model Mechanobiol 15 (2016) 1733– 1746. 10.1007/s10237-016-0794-1.

[8] C. Guillot, T. Lecuit, Mechanics of epithelial tissue homeostasis and morphogenesis, Science (1979) 340 (2013) 1185–1189. 10.1126/science.1235249.

[9] E.E. Govek, S.E. Newey, L. Van Aelst, The role of the Rho GTPases in neuronal development, Genes Dev 19 (2005) 1–49. 10.1101/gad.1256405.

[10] S. Hirano, M. Takeichi, Cadherins in brain morphogenesis and wiring, Physiol Rev 92 (2012) 597–634. 10.1152/physrev.00014.2011.

[11] V. Brault, R. Moore, S. Kutsch, M. Ishibashi, D.H. Rowitch, A.P. McMahon, L. Sommer, O. Boussadia, R. Kemler, Inactivation of the β-catenin gene by Wnt1-Cre-mediated deletion results in dramatic brain malformation and failure of craniofacial development, Development 128 (2001) 1253–1264. 10.1242/dev.128.8.1253.

[12] N. Barker, M. Huch, P. Kujala, M. van de Wetering, H.J. Snippert, J.H. van Es, T. Sato, D.E. Stange, H. Begthel, M. van den Born, E. Danenberg, S. van den Brink, J. Korving, A. Abo, P.J. Peters, N. Wright, R. Poulsom, H. Clevers, Lgr5+ve Stem Cells Drive Self-Renewal in the Stomach and Build Long-Lived Gastric Units In Vitro, Cell Stem Cell 6 (2010) 25–36. 10.1016/j.stem.2009.11.013.

[13] M. Takasato, P.X. Er, H.S. Chiu, B. Maier, G.J. Baillie, C. Ferguson, R.G. Parton, E.J. Wolvetang, M.S. Roost, S.M.C. De Sousa Lopes, M.H. Little, Kidney organoids from human iPS cells contain multiple lineages and model human nephrogenesis, Nature 526 (2015) 564–568. 10.1038/nature15695.

[14] B.R. Dye, D.R. Hill, M.A. Ferguson, Y.H. Tsai, M.S. Nagy, R. Dyal, J.M. Wells, C.N. Mayhew, R. Nattiv, O.D. Klein, E.S. White, G.H. Deutsch, J.R. Spence, In vitro generation of human pluripotent stem cell derived lung organoids, Elife 2015 (2015) e05098. 10.7554/eLife.05098.

[15] J. Lee, C.C. Rabbani, H. Gao, M.R. Steinhart, B.M. Woodru, Z.E. Pflum, A. Kim, S. Heller, Y. Liu, T.Z. Shipchandler, K.R. Koehler, Hair-bearing human skin generated entirely from pluripotent stem cells, Nature 582 (2020) 399–404. 10.1038/s41586-020-2352-3.

[16] S. Frenz-Wiessner, S.D. Fairley, M. Buser, I. Goek, K. Salewskij, G. Jonsson, D. Illig, B. zu Putlitz, D. Petersheim, Y. Li, P.H. Chen, M. Kalauz, R. Conca, M. Sterr, J. Geuder, Y. Mizoguchi, R.T.A. Megens, M.I. Linder, D. Kotlarz, M. Rudelius, J.M. Penninger, C. Marr, C. Klein, Generation of complex bone marrow organoids from human induced pluripotent stem cells, Nat Methods 21 (2024) 868–881. 10.1038/s41592-024-02172-2.

[17] M. Völkner, M. Zschätzsch, M. Rostovskaya, R.W. Overall, V. Busskamp, K. Anastassiadis, M.O. Karl, Retinal Organoids from Pluripotent Stem Cells Efficiently Recapitulate Retinogenesis, Stem Cell Reports 6 (2016) 525–538. 10.1016/j.stemcr.2016.03.001.

[18] M.A. Lancaster, M. Renner, C.A. Martin, D. Wenzel, L.S. Bicknell, M.E. Hurles, T. Homfray, J.M. Penninger, A.P. Jackson, J.A. Knoblich, Cerebral organoids model human brain development and microcephaly, Nature 501 (2013) 373–379. 10.1038/nature12517.

[19] S. Benito-Kwiecinski, S.L. Giandomenico, M. Sutclije, E.S. Riis, P. Freire-Pritchett, I. Kelava, S. Wunderlich, U. Martin, G.A. Wray, K. McDole, M.A. Lancaster, An early cell shape transition drives evolutionary expansion of the human forebrain, Cell 184 (2021) 2084–2102. 10.1016/j.cell.2021.02.050.

[20] N. Gjorevski, M. Nikolaev, T.E. Brown, O. Mitrofanova, N. Brandenberg, F.W. DelRio, F.M. Yavitt, P. Liberali, K.S. Anseth, M.P. Lutolf, Tissue geometry drives deterministic organoid patterning, Science (1979) 375 (2022) eaaw9021. 10.1126/science.aaw9021.

[21] A. Chrisnandy, D. Blondel, S. Rezakhani, N. Broguiere, M.P. Lutolf, Synthetic dynamic hydrogels promote degradation-independent in vitro organogenesis, Nat Mater 21 (2022) 479–487. 10.1038/s41563-021-01136-7.

[22] J. Zhang, D. Marciano, L. Wang, W. Wang, M. Gossen, M. Yang, T. Peng, J. Gautrot, X. Xu, N. Ma, Bioinspired Hyaluronic Acid-Based Hydrogel Fuels Bi-Directional Lung Organoid Maturation via PIEZO1 and ITGB1 Mediated Mechanosensation, Adv Mater Interfaces 11 (2024) 2400194. 10.1002/admi.202400194.

[23] M.D.A. Norman, S.A. Ferreira, G.M. Jowett, L. Bozec, E. Gentleman, Measuring the elastic modulus of soft culture surfaces and three-dimensional hydrogels using atomic force microscopy, Nat Protoc 16 (2021) 2418–2449. 10.1038/s41596-021-00495-4.

[24] O. Campàs, A toolbox to explore the mechanics of living embryonic tissues, Semin Cell Dev Biol 55 (2016) 119–130. 10.1016/j.semcdb.2016.03.011.

[25] A. Mongera, P. Rowghanian, H.J. Gustafson, E. Shelton, D.A. Kealhofer, E.K. Carn, F. Serwane, A.A. Lucio, J. Giammona, O. Campàs, A fluid-to-solid jamming transition underlies vertebrate body axis elongation, Nature 561 (2018) 401–405. 10.1038/s41586-018-0479-2.

[26] O. Campàs, T. Mammoto, S. Hasso, R.A. Sperling, D. O’connell, A.G. Bischof, R. Maas, D.A. Weitz, L. Mahadevan, D.E. Ingber, Quantifying cell-generated mechanical forces within living embryonic tissues, Nat Methods 11 (2014) 183–189. 10.1038/nmeth.2761.

[27] N.P. Shro, P. Xu, S. Kim, E.R. Shelton, B.J. Gross, Y. Liu, C.O. Gomez, Q. Ye, T.Y. Drennon, J.K. Hu, J.B.A. Green, O. Campàs, O.D. Klein, Proliferation-driven mechanical compression induces signalling centre formation during mammalian organ development, Nat Cell Biol 26 (2024) 519–529. 10.1038/s41556-024-01380-4.

[28] S. Dupont, L. Morsut, M. Aragona, E. Enzo, S. Giulitti, M. Cordenonsi, F. Zanconato, J. Le Digabel, M. Forcato, S. Bicciato, N. Elvassore, S. Piccolo, Role of YAP/TAZ in mechanotransduction, Nature 474 (2011) 179–183. 10.1038/nature10137.

[29] G. Brusatin, T. Panciera, A. Gandin, A. Citron, S. Piccolo, Biomaterials and engineered microenvironments to control YAP/TAZ-dependent cell behaviour, Nat Mater 17 (2018) 1063–1075. 10.1038/s41563-018-0180-8.

[30] B.K. Terry, S. Kim, The role of Hippo-YAP/TAZ signaling in brain development, Developmental Dynamics 251 (2022) 1644–1665. 10.1002/dvdy.504.

[31] J. Wang, Y. Xiao, C.W. Hsu, I.M. Martinez-Traverso, M. Zhang, Y. Bai, M. Ishii, R.E. Maxson, E.N. Olson, M.E. Dickinson, J.D. Wythe, J.F. Martin, Yap and taz play a crucial role in neural crest-derived craniofacial development, Development (Cambridge) 143 (2016) 504–515. 10.1242/dev.126920.

[32] C.J. Hindley, A.L. Condurat, V. Menon, R. Thomas, L.M. Azmitia, J.A. Davis, J. Pruszak, The Hippo pathway member YAP enhances human neural crest cell fate and migration, Sci Rep 6 (2016) 23208. 10.1038/srep23208.

[33] X. Cao, S.L. Pfa, F.H. Gage, YAP regulates neural progenitor cell number via the TEA domain transcription factor, Genes Dev 22 (2008) 3320–3334. 10.1101/gad.1726608.

[34] A. Lavado, R. Gangwar, J. Paré, S. Wan, Y. Fan, X. Cao, YAP/TAZ maintain the proliferative capacity and structural organization of radial glial cells during brain development, Dev Biol 480 (2021) 39–49. 10.1016/j.ydbio.2021.08.010.

[35] W.A. Liu, S. Chen, Z. Li, C.H. Lee, G. Mirzaa, W.B. Dobyns, M.E. Ross, J. Zhang, S.H. Shi, PARD3 dysfunction in conjunction with dynamic HIPPO signaling drives cortical enlargement with massive heterotopia, Genes Dev 32 (2018) 763–780. 10.1101/gad.313171.118.

[36] L.J. Hughes, R. Park, M.J. Lee, B.K. Terry, D.J. Lee, H. Kim, S.H. Cho, S. Kim, Yap/Taz are required for establishing the cerebellar radial glia scaffold and proper foliation, Dev Biol 457 (2020) 150–162. 10.1016/j.ydbio.2019.10.002.

[37] J.Y. Kim, R. Park, J.H.J. Lee, J. Shin, J. Nickas, S. Kim, S.H. Cho, Yap is essential for retinal progenitor cell cycle progression and RPE cell fate acquisition in the developing mouse eye, Dev Biol 419 (2016) 336–347. 10.1016/j.ydbio.2016.09.001.

[38] B.V.V.G. Reddy, C. Rauskolb, K.D. Irvine, Influence of Fat-Hippo and Notch signaling on the proliferation and differentiation of Drosophila optic neuroepithelia, Development 137 (2010) 2397–2408. 10.1242/dev.050013.

[39] Z. Meng, Y. Qiu, K.C. Lin, A. Kumar, J.K. Placone, C. Fang, K.C. Wang, S. Lu, M. Pan, A.W. Hong, T. Moroishi, M. Luo, S.W. Ploue, Y. Diao, Z. Ye, H.W. Park, X. Wang, F.X. Yu, S. Chien, C.Y. Wang, B. Ren, A.J. Engler, K.L. Guan, RAP2 mediates mechanoresponses of the Hippo pathway, Nature 560 (2018) 655–660. 10.1038/s41586-018-0444-0.

[40] A. Das, R.S. Fischer, D. Pan, C.M. Waterman, YAP nuclear localization in the absence of cell-cell contact is mediated by a filamentous actin-dependent, Myosin IIand Phospho-YAP-independent pathway during extracellular matrix mechanosensing, Journal of Biological Chemistry 291 (2016) 6096–6110. 10.1074/jbc.M115.708313.

[41] G. Nardone, J. Oliver-De La Cruz, J. Vrbsky, C. Martini, J. Pribyl, P. Skládal, M. Pešl, G. Caluori, S. Pagliari, F. Martino, Z. Maceckova, M. Hajduch, A. Sanz-Garcia, N.M. Pugno, G.B. Stokin, G. Forte, YAP regulates cell mechanics by controlling focal adhesion assembly, Nat Commun 8 (2017) 15321. 10.1038/ncomms15321.

[42] C. Loebel, R.L. Mauck, J.A. Burdick, Local nascent protein deposition and remodelling guide mesenchymal stromal cell mechanosensing and fate in three-dimensional hydrogels, Nat Mater 18 (2019) 883–891. 10.1038/s41563-019-0307-6.

[43] J. Oliver-De La Cruz, G. Nardone, J. Vrbsky, A. Pompeiano, A.R. Perestrelo, F. Capradossi, K. Melajová, P. Filipensky, G. Forte, Substrate mechanics controls adipogenesis through YAP phosphorylation by dictating cell spreading, Biomaterials 205 (2019) 64–80. 10.1016/j.biomaterials.2019.03.009.

[44] M. Fischer, P. Rikeit, P. Knaus, C. Coirault, YAP-mediated mechanotransduction in skeletal muscle, Front Physiol 7 (2016) 41. 10.3389/fphys.2016.00041.

[45] T. Heallen, M. Zhang, J. Wang, M. Bonilla-Claudio, E. Klysik, R.L. Johnson, J.F. Martin, Hippo pathway inhibits wnt signaling to restrain cardiomyocyte proliferation and heart size, Science (1979) 332 (2011) 458–461. 10.1126/science.1199010.

[46] S.J.I. Blackford, T.T.L. Yu, M.D.A. Norman, A.M. Syanda, M. Manolakakis, D. Lachowski, Z. Yan, Y. Guo, E. Garitta, F. Riccio, G.M. Jowett, S.S. Ng, S. Vernia, A.E. del Río Hernández, E. Gentleman, S.T. Rashid, RGD density along with substrate stiffness regulate hPSC hepatocyte functionality through YAP signalling, Biomaterials 293 (2023) 121982. 10.1016/j.biomaterials.2022.121982.

[47] H. Zhang, H.A. Pasolli, E. Fuchs, Yes-associated protein (YAP) transcriptional coactivator functions in balancing growth and differentiation in skin, Proc Natl Acad Sci U S A 108 (2011) 2270–2275. 10.1073/pnas.1019603108.

[48] G. Walko, S. Woodhouse, A.O. Pisco, E. Rognoni, K. Liakath-Ali, B.M. Lichtenberger, A. Mishra, S.B. Telerman, P. Viswanathan, M. Logtenberg, L.M. Renz, G. Donati, S.R. Quist, F.M. Watt, A genome-wide screen identifies YAP/WBP2 interplay conferring growth advantage on human epidermal stem cells, Nat Commun 8 (2017) 14744. 10.1038/ncomms14744.

[49] D. Pinheiro, Y. Bellaïche, Mechanical Force-Driven Adherens Junction Remodeling and Epithelial Dynamics, Dev Cell 47 (2018) 3–19. 10.1016/j.devcel.2018.09.014.

[50] B.D. Cosgrove, K.L. Mui, T.P. Driscoll, S.R. Caliari, K.D. Mehta, R.K. Assoian, J.A. Burdick, R.L. Mauck, N-cadherin adhesive interactions modulate matrix mechanosensing and fate commitment of mesenchymal stem cells, Nat Mater 15 (2016) 1297–1306. 10.1038/nmat4725.

[51] D. Serra, U. Mayr, A. Boni, I. Lukonin, M. Rempfler, L. Challet Meylan, M.B. Stadler, P. Strnad, P. Papasaikas, D. Vischi, A. Waldt, G. Roma, P. Liberali, Self-organization and symmetry breaking in intestinal organoid development, Nature 569 (2019) 66–72. 10.1038/s41586-019-1146-y.

[52] N. Gjorevski, N. Sachs, A. Manfrin, S. Giger, M.E. Bragina, P. Ordóñez-Morán, H. Clevers, M.P. Lutolf, Designer matrices for intestinal stem cell and organoid culture, Nature 539 (2016) 560–564. 10.1038/nature20168.

[53] D. Yimlamai, C. Christodoulou, G.G. Galli, K. Yanger, B. Pepe-Mooney, B. Gurung, K. Shrestha, P. Cahan, B.Z. Stanger, F.D. Camargo, Hippo pathway activity influences liver cell fate, Cell 157 (2014) 1324–1338. 10.1016/j.cell.2014.03.060.

[54] G. Sorrentino, S. Rezakhani, E. Yildiz, S. Nuciforo, M.H. Heim, M.P. Lutolf, K. Schoonjans, Mechano-modulatory synthetic niches for liver organoid derivation, Nat Commun 11 (2020) 3416. 10.1038/s41467-020-17161-0.

[55] Q. Tan, K.M. Choi, D. Sicard, D.J. Tschumperlin, Human airway organoid engineering as a step toward lung regeneration and disease modeling, Biomaterials 113 (2017) 118–132. 10.1016/j.biomaterials.2016.10.046.

[56] X.-H. Li, D. Guo, L.-Q. Chen, Z.-H. Chang, J.-X. Shi, N. Hu, C. Chen, X.-W. Zhang, S.-Q. Bao, M.-M. Chen, D. Ming, Low-intensity ultrasound ameliorates brain organoid integration and rescues microcephaly deficits, Brain 147 (2024) 3817– 3833. 10.1093/brain/awae150.

[57] D. Guo, B. Yao, W. Shao, J. Zuo, Z. Chang, J. Shi, N. Hu, S. Bao, M. Chen, X. Fan, X. Li, The Critical Role of YAP/BMP/ID1 Axis on Simulated Microgravity-Induced Neural Tube Defects in Human Brain Organoids, Advanced Science 12 (2025) 2410188. 10.1002/advs.202410188.

[58] C. Tang, X. Wang, M. D’Urso, C. van der Putten, N.A. Kurniawan, 3D Interfacial and Spatiotemporal Regulation of Human Neuroepithelial Organoids, Advanced Science 9 (2022) 2201106. 10.1002/advs.202201106.

[59] J. Ha, J.S. Kang, M. Lee, A. Baek, S. Kim, S.K. Chung, M.O. Lee, J. Kim, Simplified Brain Organoids for Rapid and Robust Modeling of Brain Disease, Front Cell Dev Biol 8 (2020) 594090. 10.3389/fcell.2020.594090.

[60] S. Goswami, I. Balasubramanian, L. D’Agostino, S. Bandyopadhyay, R. Patel, S. Avasthi, S. Yu, J.R. Goldenring, E.M. Bonder, N. Gao, RAB11A-mediated YAP localization to adherens and tightjunctions is essential for colonic epithelial integrity, Journal of Biological Chemistry 297 (2021) 100848. 10.1016/j.jbc.2021.100848.

[61] G.C. Fletcher, M. del C. Diaz-de-la-Loza, N. Borreguero-Muñoz, M. Holder, M. Aguilar-Aragon, B.J. Thompson, Mechanical strain regulates the Hippo pathway in Drosophila, Development (Cambridge) 145 (2018) 159467. 10.1242/dev.159467.

[62] H. Hirata, M. Samsonov, M. Sokabe, Actomyosin contractility provokes contact inhibition in E-cadherin-ligated keratinocytes, Sci Rep 7 (2017) 46326. 10.1038/srep46326.

[63] K.T. Furukawa, K. Yamashita, N. Sakurai, S. Ohno, The Epithelial Circumferential Actin Belt Regulates YAP/TAZ through Nucleocytoplasmic Shuttling of Merlin, Cell Rep 20 (2017) 1435–1447. 10.1016/j.celrep.2017.07.032.

[64] D. Kong, L. Peng, M. Bosch-Fortea, A. Chrysanthou, C.V.J.-M. Alexis, C. Matellan, A. Zarbakhsh, G. Mastroianni, A. del Rio Hernandez, J.E. Gautrot, Impact of the multiscale viscoelasticity of quasi-2D self-assembled protein networks on stem cell expansion at liquid interfaces, Biomaterials 284 (2022) 121494. 10.1016/j.biomaterials.2022.121494.

[65] A. Chrysanthou, M. Bosch-Fortea, J.E. Gautrot, Co-Surfactant-Free Bioactive Protein Nanosheets for the Stabilization of Bioemulsions Enabling Adherent Cell Expansion, Biomacromolecules 24 (2023) 4465–4477. 10.1021/acs.biomac.2c01289.

[66] D. Kong, L. Peng, S. Di Cio, P. Novak, J.E. Gautrot, Stem Cell Expansion and Fate Decision on Liquid Substrates Are Regulated by Self-Assembled Nanosheets, ACS Nano 12 (2018) 9206–9213. 10.1021/acsnano.8b03865.

[67] D. Kong, W. Megone, K.D.Q. Nguyen, S. Di Cio, M. Ramstedt, J.E. Gautrot, Protein Nanosheet Mechanics Controls Cell Adhesion and Expansion on Low-Viscosity Liquids, Nano Lett 18 (2018) 1946–1951. 10.1021/acs.nanolett.7b05339.

[68] L. Peng, J.E. Gautrot, Long term expansion profile of mesenchymal stromal cells at protein nanosheet-stabilised bioemulsions for next generation cell culture microcarriers, Mater Today Bio 12 (2021) 100159. 10.1016/j.mtbio.2021.100159.

[69] X. Jia, K. Minami, K. Uto, A.C. Chang, J.P. Hill, J. Nakanishi, K. Ariga, Adaptive Liquid Interfacially Assembled Protein Nanosheets for Guiding Mesenchymal Stem Cell Fate, Advanced Materials 32 (2020) 1905942. 10.1002/adma.201905942.

[70] X. Jia, J. Song, W. Lv, J.P. Hill, J. Nakanishi, K. Ariga, Adaptive liquid interfaces induce neuronal differentiation of mesenchymal stem cells through lipid raft assembly, Nat Commun 13 (2022) 3110. 10.1038/s41467-022-30622-y.

[71] E. Mojares, C. Nadal, D. Hayler, H. Kanso, A. Chrysanthou, C. Neri Cruz, J. Gautrot, Strong Elastic Protein Nanosheets Enable the Culture and Differentiation of Induced Pluripotent Stem Cells on Microdroplets, Adv. Mater. 36 (2024) 2406333. 10.1002/adma.202406333.

[72] J. Gautrot, D. Marciano, M. Bosch-Fortea, Biomimetic Artificial Bone Marrow Niches for the Scale Up of Hematopoietic Stem Cells, BioRxiv (2023). doi10.1101/2023.06.23.546233.

[73] A. Chrysanthou, M. Bosch-Fortea, C. Nadal, A. Zarbakhsh, J. Gautrot, Interfacial Mechanics of b-Casein and Albumin Mixed Protein Assemblies at Liquid-Liquid Interfaces, J. Colloid Interface Sci. 674 (2024) 379–391.

[74] C. Mücksch, H.M. Urbassek, Molecular dynamics simulation of free and forced BSA adsorption on a hydrophobic graphite surface, Langmuir 27 (2011) 12938– 12943. 10.1021/la201972f.

[75] J.R. Simard, P.A. Zunszain, C.E. Ha, J.S. Yang, N. V. Bhagavan, I. Petitpas, S. Curry, J.A. Hamilton, Locating high-affinity fatty acid-binding sites on albumin by x-ray crystallography and NMR spectroscopy, Proc Natl Acad Sci U S A 102 (2005) 17958–17963. 10.1073/pnas.0506440102.

[76] J. Jumper, R. Evans, A. Pritzel, T. Green, M. Figurnov, O. Ronneberger, K. Tunyasuvunakool, R. Bates, A. Žídek, A. Potapenko, A. Bridgland, C. Meyer, S.A.A. Kohl, A.J. Ballard, A. Cowie, B. Romera-Paredes, S. Nikolov, R. Jain, J. Adler, T. Back, S. Petersen, D. Reiman, E. Clancy, M. Zielinski, M. Steinegger, M. Pacholska, T. Berghammer, S. Bodenstein, D. Silver, O. Vinyals, A.W. Senior, K. Kavukcuoglu, P. Kohli, D. Hassabis, Highly accurate protein structure prediction with AlphaFold, Nature 596 (2021) 583–589. 10.1038/s41586-021-03819-2.

[77] W. Zhu, G. Gong, J. Pan, S. Han, W. Zhang, Y. Hu, L. Xie, High level expression and purification of recombinant human serum albumin in Pichia pastoris, Protein Expr Purif 147 (2018) 61–68. 10.1016/j.pep.2018.02.003.

[78] G.P. Lin-Cereghino, C.M. Stark, D. Kim, J. Chang, N. Shaheen, H. Poerwanto, K. Agari, P. Moua, L.K. Low, N. Tran, A.D. Huang, M. Nattestad, K.T. Oshiro, J.W. Chang, A. Chavan, J.W. Tsai, J. Lin-Cereghino, The effect of α-mating factor secretion signal mutations on recombinant protein expression in Pichia pastoris, Gene 519 (2013) 311–317. 10.1016/j.gene.2013.01.062.

[79] J. Yu, Y. Chen, L. Xiong, X. Zhang, Y. Zheng, Conductance changes in Bovine Serum Albumin caused by drug-binding triggered structural transitions, Materials 12 (2019) 1022. 10.3390/ma12071022.

[80] A. Chrysanthou, H. Kanso, W. Zhong, L. Shang, J.E. Gautrot, Supercharged Protein Nanosheets for Cell Expansion on Bioemulsions, ACS Appl Mater Interfaces 15 (2023) 2760–2770. 10.1021/acsami.2c20188.

[81] E.M. Freer, K.S. Yim, G.G. Fuller, C.J. Radke, Interfacial Rheology of Globular and Flexible Proteins at the Hexadecane/Water Interface: Comparison of Shear and Dilatation Deformation, Journal of Physical Chemistry B 108 (2004). 10.1021/jp037236k.

[82] Y.S. Hwang, G.C. Bong, D. Ortmann, N. Hattori, H.C. Moeller, A. Khademhosseinia, Microwell-mediated control of embryoid body size regulates embryonic stem cell fate via differential expression of WNT5a and WNT11, Proc Natl Acad Sci U S A 106 (2009) 16978–16983. 10.1073/pnas.0905550106.

[83] S. Novosedlik, F. Reichel, T. van Veldhuisen, Y. Li, H. Wu, H. Janssen, J. Guck, J. van Hest, Cytoskeleton-functionalized synthetic cells with life-like mechanical features and regulated membrane dynamicity, Nat Chem 17 (2025) 356–364. 10.1038/s41557-024-01697-5.

[84] M. Abbas, W.P. Lipiński, K.K. Nakashima, W.T.S. Huck, E. Spruijt, A short peptide synthon for liquid–liquid phase separation, Nat Chem 13 (2021) 1046–1054. 10.1038/s41557-021-00788-x.

[85] O. Rifaie-Graham, J. Yeow, A. Najer, R. Wang, R. Sun, K. Zhou, T.N. Dell, C. Adrianus, C. Thanapongpibul, M. Chami, S. Mann, J.R. de Alaniz, M.M. Stevens, Photoswitchable gating of non-equilibrium enzymatic feedback in chemically communicating polymersome nanoreactors, Nat Chem 15 (2023) 110–118. 10.1038/s41557-022-01062-4.

[86] H. Kim, J. Yeow, A. Najer, W. Kit-Anan, R. Wang, O. Rifaie-Graham, C. Thanapongpibul, M.M. Stevens, Microliter Scale Synthesis of Luciferase-Encapsulated Polymersomes as Artificial Organelles for Optogenetic Modulation of Cardiomyocyte Beating, Advanced Science 9 (2022) 2200239. 10.1002/advs.202200239.

[87] S.R. Caliari, J.A. Burdick, A practical guide to hydrogels for cell culture, Nat Methods 13 (2016) 405–414. 10.1038/nmeth.3839.

