## Supplementary Information for "Artificial Cells Evidence Apical Compressive Forces Building up During Neuroepithelial Organoid Early Development"

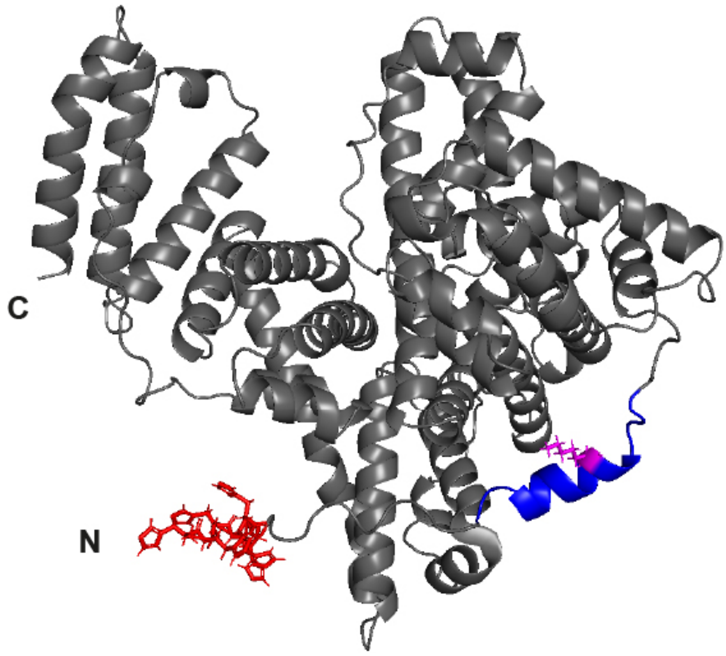


**Supplementary Figure S1.** Pymol representation of the predicted secondary structure for rBSA-Avi, generated by Alpha Fold 2. The N- and C- terminal regions are indicated by corresponding letters. The His6-tag is indicated in red, and the Avi-tag in blue, with the lysine residue within the Avi-tag (biotinylated by rBirA enzyme) indicated in pink.

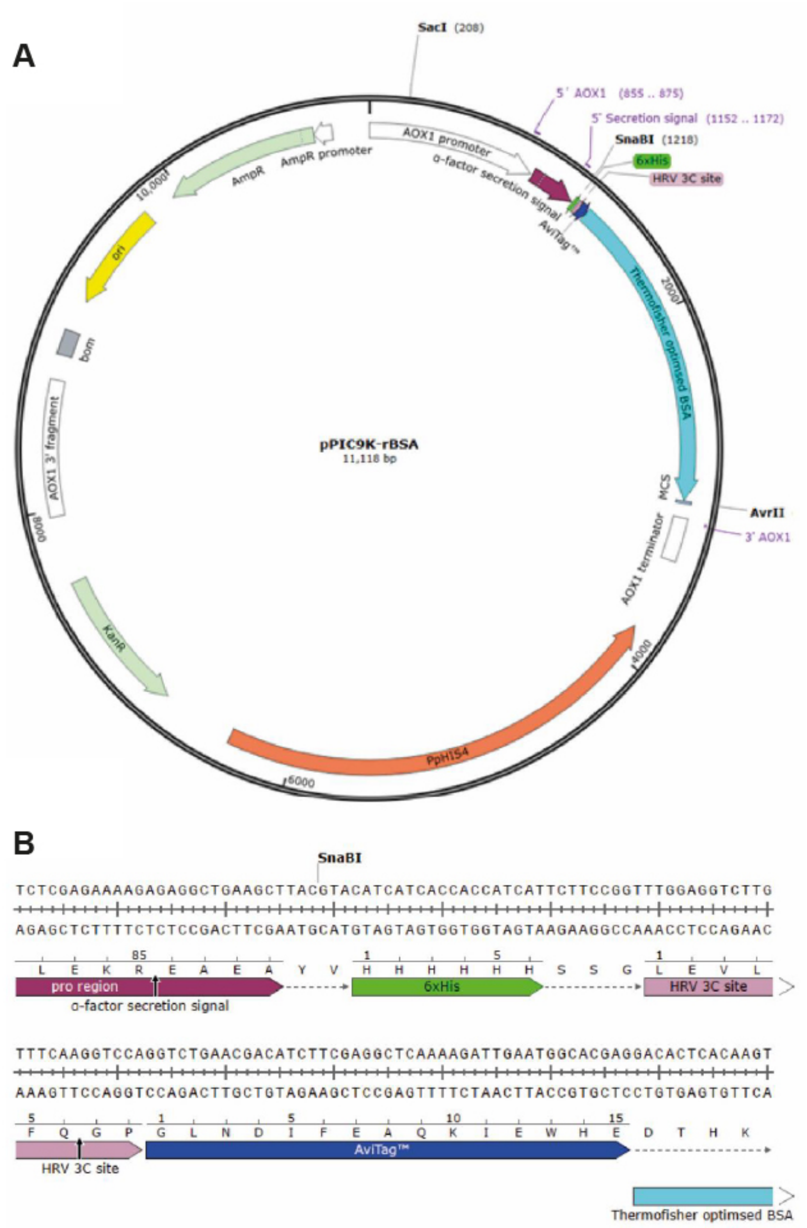


**Supplementary Figure S2.** P. pastoris expression vector for Avi-rBSA. A. Schematic showing the pPIC9K-Avi-rBSA plasmid constructed using the SnapGene software. Light green: KanR = Geneticin (G418) resistance, AmpR = Ampicillin resistance. Cyan = Avi-rBSA insert optimised using the ThermoFisher GeneArt service. Maroon: Alpha-factor secretion signal. White = AOX1 promoter and fragments. Green = 6x His-Tag. Blue = Avi-Tag. Pink = HRV 3C cleavage site. Highlighted are the restriction enzyme cleavage sites AvrII and SnaBI. B. Magnified schematic showing the detailed N-terminal region of the pPIC9K-Avi-rBSA sequence constructed using the SnapGene software. Cyan = Avi-rBSA insert optimised using the ThermoFisher GeneArt service. Maroon: Alpha-factor secretion signal. White = AOX1 promoter. Green = 6x His-Tag. Pink = Avi-Tag. Light Pink = HRV 3C cleavage site.

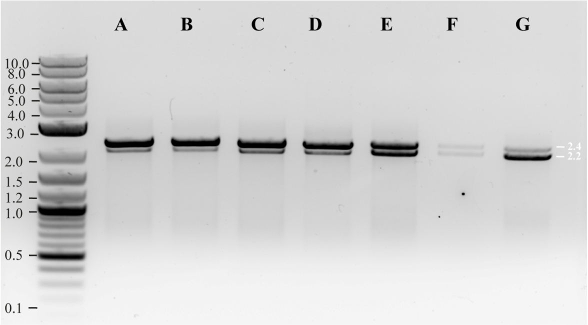


**Supplementary Figure S3.** PCR of isolated chromosomal DNA for GS115 Pichia pastoris transformants that were plated onto varying concentrations of G418 post His+ selection, run on a 0.8% agarose gel. Primers: 5' AOX1 + 3' AOX1. Positive transformants exhibit two bands, one corresponding to the AOX1 promoter in the Pichia pastoris genome (2.2 kbp) and a second for the inserted recombinant BSA sequence (2.45 kbp). The bands are: A-B) 4.0 mg/mL G418 isolated colonies. C-D) 2.0 mg/mL G418 isolated colonies. E) 1.0 mg/mL G418 isolated colonies. F) 0.5 mg/mL G418 isolated colonies. G) 0.25 mg/mL G418 isolated colonies. DNA ladder is in kbp.

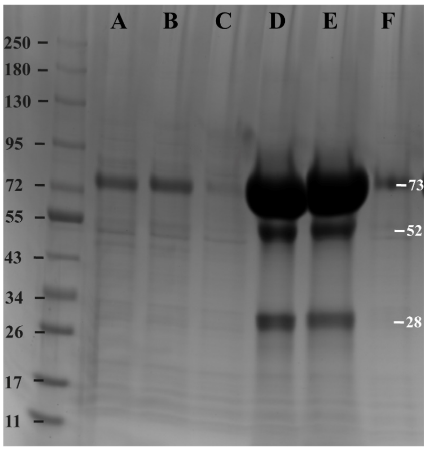


**Supplementary Figure S4.** SDS page gel confirming the recombinant expression and purification of rBSA-Avi in P. pastoris, induced with 2% methanol for 4 days at 28$^{o}$C. Purification was with his-tag nickel affinity chromatography, column washing using PBS + 20 mM imidazole, followed by elution using two in tandem HisTrap columns with PBS + 500 mM imidazole. The bands are: A) Sample collected after 4 day expression with 2% methanol at 28$^{o}$C. B) Loading fraction. C) Wash fraction. D-E) Elution fractions 11-17. F) Flow Through fraction. Protein ladder is in kDa.

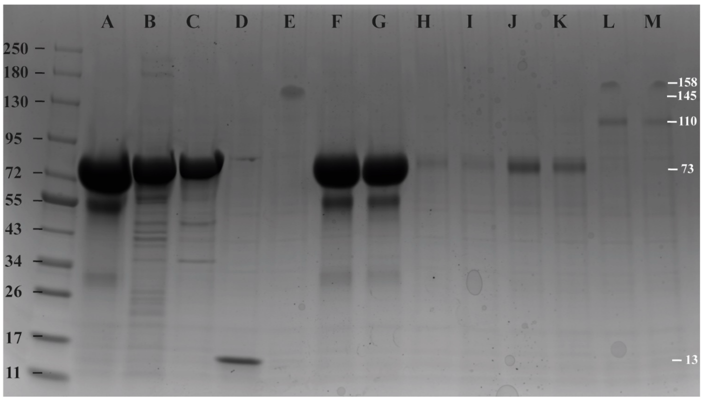


**Supplementary Figure S5.** 4-20% SDS page gel confirming the successful biotinylation of rBSA-Avi (at 45 μM), using 3 μM rBirA and 0.3 mM D-Biotin. rBSA-Avi runs at approximately 71 kDa. rBirA-MBP runs at approximately 78 kDa. Native Streptavidin runs at 150 kDa. Streptavinated-rBSA-Bt complexes run at approximately 110 kDa. The bands are: A) rBSA-Avi. B-C) rBirA-MBP. D) 1 mg/mL Streptavidin boiled at 95°C. E) 1 mg/mL Streptavidin native. F-G) rBSA-Bt. H-I) rBSA-Bt post-PD10 column fractions. J-K) rBSA-Bt post 30K MWCO concentrator fractions. L-M) rBSA-Bt with the addition of 1mg/mL native Streptavidin. The Protein ladder is in kDa.

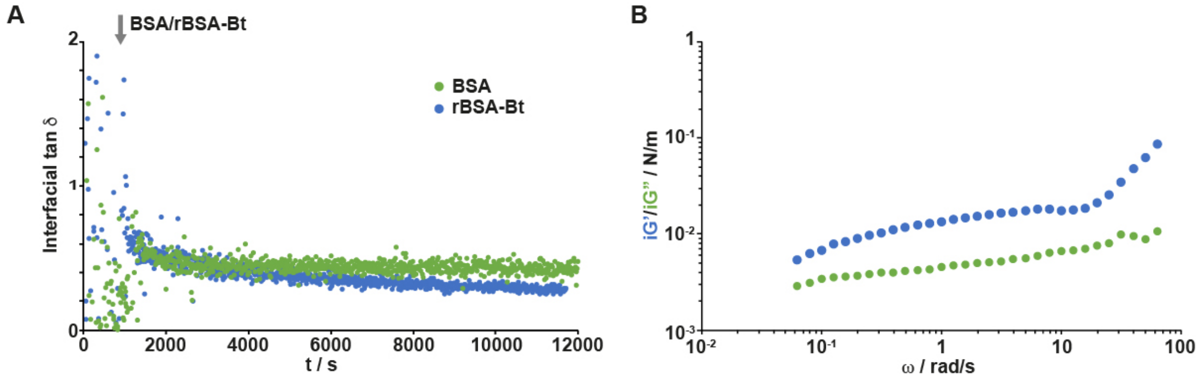


**Supplementary Figure S6.** Representative time sweeps (frequency, 0.1 Hz; strain, $10^{-3}$ rad) monitoring the evolution of the interfacial tan δ during the assembly of rBSA-Bt and BSA at the surface of Novec 7500 oil, in PBS (protein concentration: 1 mg/mL). The arrow indicates the time point of injection of corresponding proteins. B. Frequency sweeps of interfaces generated by sequential assembly of rBSA-Bt and streptavidin (strain of $10^{-3}$ rad).

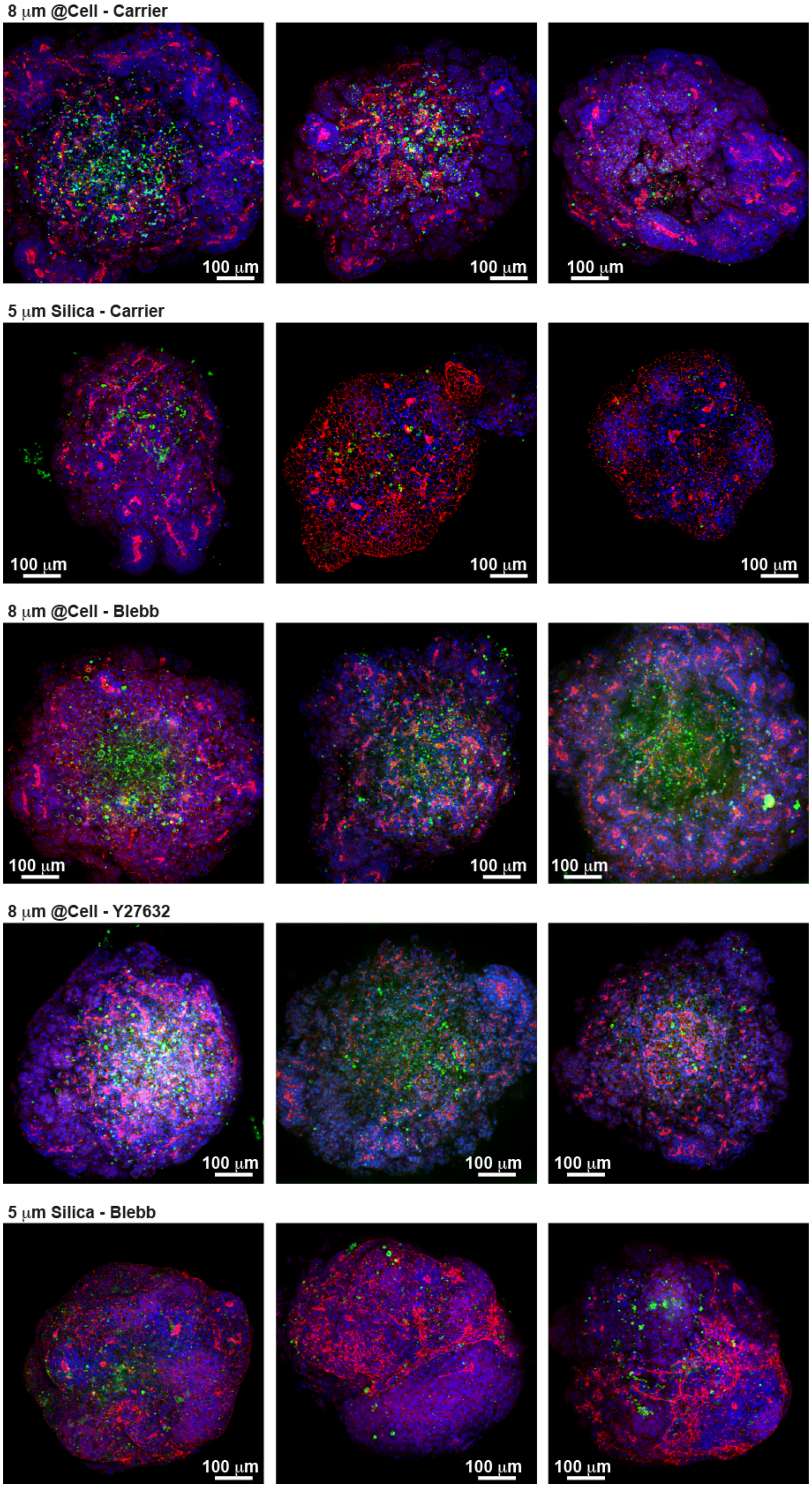


**Supplementary Figure S7.** A. Confocal microscopy images (Max projections of z-stacks; 3 representative organoids shown per condition) of neuroepithelial organoids seeded with 8 μm artificial cells (@Cell) or 5 μm silica microparticles presenting E/N-cadherins, at day 3 post induction, with and without Blebbistatin/Y27632 (10 and 20 μM, respectively; Blue, DAPI; Green, Streptavidin; Red, ZO1).

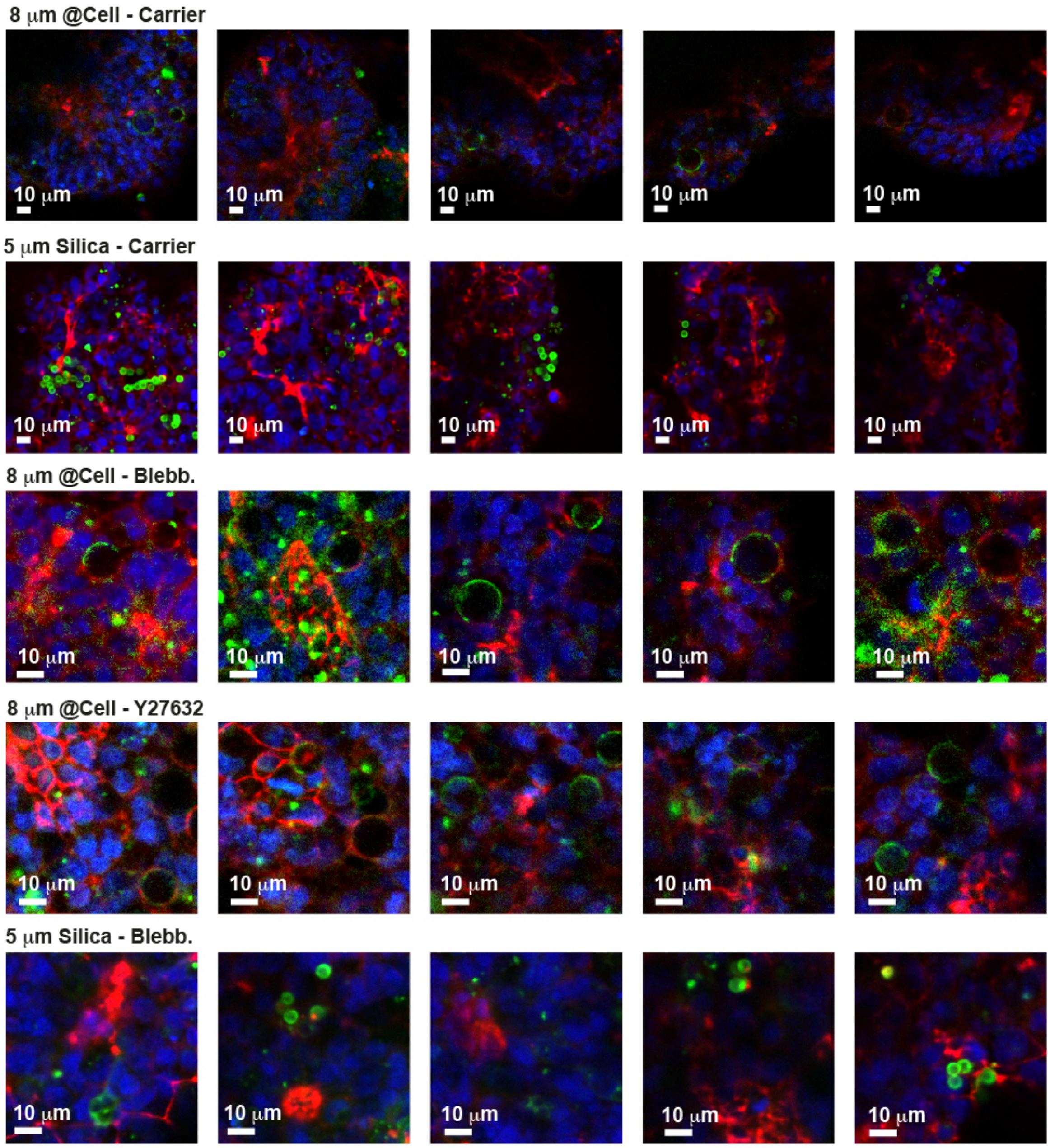


**Supplementary Figure S8.** Example of zooms taken from individual slices, from confocal stacks organoids seeded with artificial cells or silica microparticles presenting E/N-cadherins, at day 3 post induction, with and without Blebbistatin/Y27632 (10 and 20 μM, respectively; Blue, DAPI; Green, Streptavidin; Red, ZO1). 5 representative images shown for each condition.

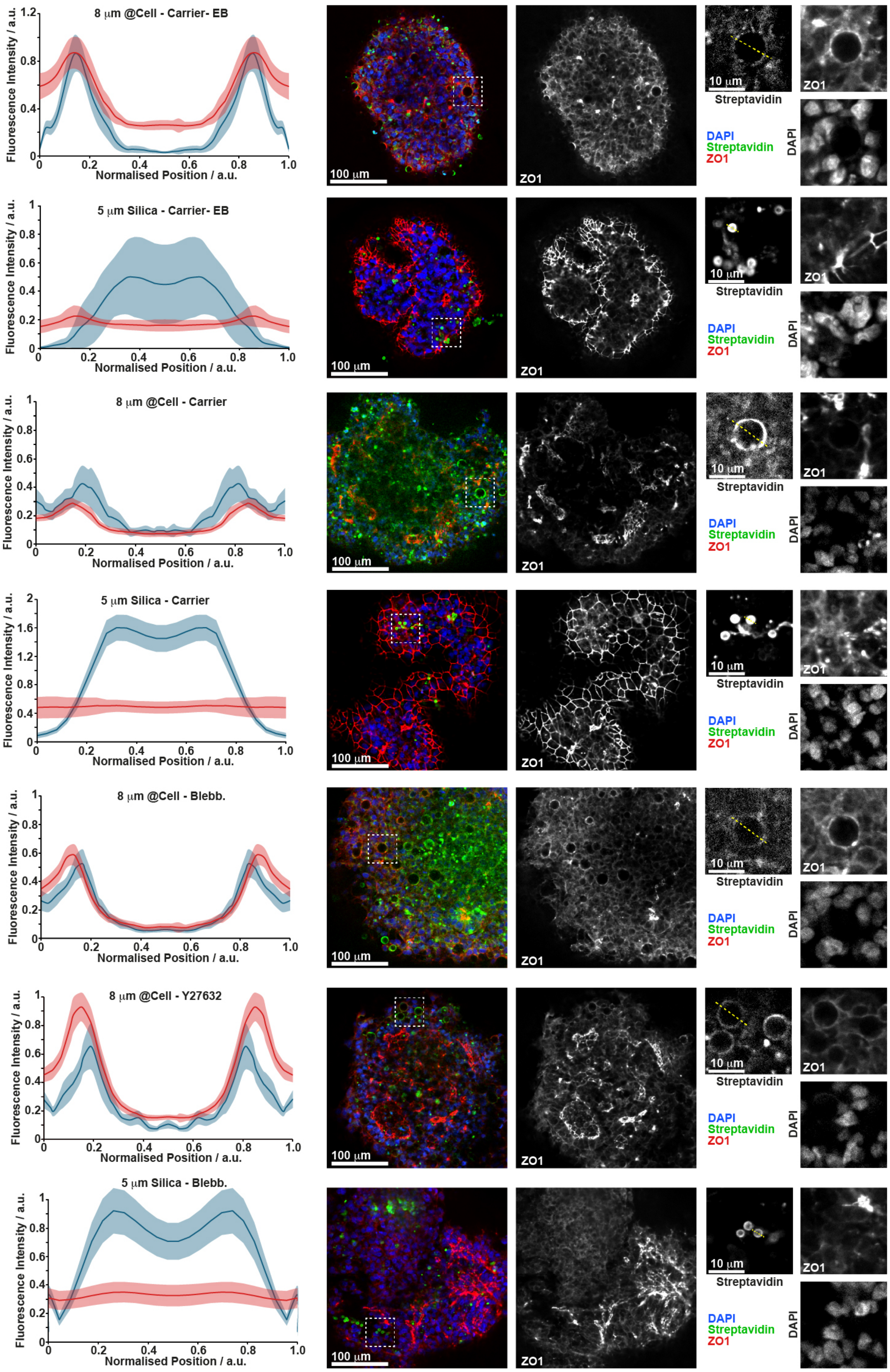
8 μm @Cell - Carrier- EB
5 μm Silica - Carrier- EB
8 μm @Cell - Carrier
5 μm Silica - Carrier
8 μm @Cell - Blebb.
8 μm @Cell - Y27632
5 μm Silica - Blebb.
Fluorescence Intensity / a.u.
Normalised Position / a.u.
100 μm
10 μm
ZO1
Streptavidin
DAPI
Streptavidin
ZO1
DAPI

**Supplementary Figure S9.** Examples of single optical slices taken from stacks and zooms from embryoid bodies (EB) and neuroepithelial organoids cultured for 3 days post induction, showing individual channels (Blue, DAPI; Green, Streptavidin; Red, ZO1), together with the quantification of streptavidin (blue) and ZO1(red) intensity profiles (shaded areas correspond to associated standard deviations from >10 profiles).

#### Supplementary Videos

**Supplementary Video S1.** Video of neuroepithelial organoid development (days 2-3 post induction).

**Supplementary Video S2.** Video of neuroepithelial organoid development (days 3-5 post induction).

**Supplementary Video S3.** Video of embryoid body formation in the presence of artificial cells (functionalised with E-cadherin).

**Supplementary Video S4.** Video of artificial-seeded neuroepithelial organoid development (days 2-3 post induction).

### Supplementary Tables

**Supplementary Table S1: Digestion master mix for the digestion of pEX-A2-BSA and pPIC9K plasmids. 16 µL of each plasmid was digested in separate Eppendorf tubes. Total reaction volume is 20 µL.**

| | pEX-A2-rBSA | pPIC9K |
|---|---|---|
| **10x FD Green buffer** | 2 µL | 1 µL |
| **AvrII endonuclease** | 1 µL | 0.5 µL |
| **SnaBI endonuclease** | 1 µL | 0.5 µL |

**Supplementary Table S2: Digestion master mix for the analytical digest of miniprepped pPIC9K-rBSA plasmids.** SnaBI recognizes TAC^GTA sites. AvrII recognizes C^CTAGG sites.

| **Component** | **Volume (µL)** |
|---|---|
| 10x FastDigest Green buffer | 1 |
| SnaBI endonuclease | 0.5 |
| AvrII endonuclease | 0.5 |
| $ddH_2O$ | 5 |
| pPIC9K-rBSA | 3 |

**Supplementary Table S3. Digestion mix used to linearise the pPIC9K-rBSA plasmid using SacI restriction enzyme; recognizes GAGCT^C sites.**

| **Component** | **Volume (µL)** |
|---|---|
| pPIC9K-rBSA (~200ng/µL) | 5 |
| SacI Fast digest | 1 |
| $ddH_2O$ | 12 |
| Fast digest Buffer | 2 |

**Supplementary Table S4: The 5' and 3' AOX1, and the Alpha secretion signal primers**

| | |
|---|---|
| 5' AOX1 | GACTGGTTCCAATTGACAAGC |
| 3' AOX1 | GCAAATGGCATTCTGACATCC |
| Alpha secretion signal | TACTATTGCCAGCATTGCTGC |

**Supplementary Table S5: The components used to generate the master mix for colony PCR samples.**

| Components | Volume (µL) |
|---|---|
| OneTaq® 2X Master Mix | 10 |
| 10µM 5' AOX1 Primer | 0.5 |
| 10µM 3' AOX1 Primer | 0.5 |
| $ddH_2O$ | 7 |
| DNA | 2 |

**Supplementary Table S6: The components used for the invitro biotinylation of Avi-BSA.**

| Components | Final concentration |
|---|---|
| Avi-Tagged protein in PBS | 175 µM |
| rBirA | 3 µM |
| Magnesium Chloride | 10 mM |
| ATP | 10 mM |
| D-Biotin | 0.3 mM |

**Supplementary Table S7: Media used for the thawing, passaging and maintenance of hIPSCs.**

| Medium | Component | Volumes |
|---|---|---|
| E8 Flex complete | Essential 8 Flex Basal Medium | 24.5 mL |
| | Essential 8 Flex Supplement | 500 µL |
| E8 Seeding | E8 flex complete | 9990 µL |
| (Per aliquot) | 10mM Rock inhibitor (Y-27632) | 10 µL (10 µM) |
| Cell detachment solution | 0.5M EDTA (pH 8.0) | 1 µL |
| (Per well) | dPBS | 999 µL |
| Matrigel solution | DMEM/F12 | 5700 µL |
| (Per 6 wells) | Matrigel® (GFR) | 300 µL |

**Supplementary Table S8: Media used for the generation of 30 embryoid bodies.**

| Medium | Component | Volumes |
|---|---|---|
| EB formation | STEMdiff™ Cerebral Organoid Basal Medium 1 | 8 mL |
| | STEMdiff™ Cerebral Organoid Supplement A | 2 mL |
| EB seeding | EB formation medium | 6994 µL |
| | 10mM Rock inhibitor (Y-27632) | 7 µL (10 µM) |
| Dissociation | Gentle Cell Dissociation Reagent | 1 mL |

| Supplementary Table S9: Media used for the generation of cerebral organoids. | | |
|---|---|---|
| **Medium** | **Component** | **Volumes** |
| Expansion | STEMdiff™ Cerebral Organoid Basal Medium 2 | 5,700 µL |
| | STEMdiff™ Cerebral Organoid Supplement C | 60 µL |
| | STEMdiff™ Cerebral Organoid Supplement D | 120 µL |
| | Matrigel for culture (2% Final) | 120 µL |
| Maturation | STEMdiff™ Cerebral Organoid Basal Medium 2 | 5,760 µL |
| | STEMdiff™ Cerebral Organoid Supplement E | 120 µL |
| | Matrigel for culture (2% Final in solution) | 120 µL |
